# Eukaryotic secreted proteins are encoded in repeat-rich genomic regions

**DOI:** 10.64898/2026.03.17.712334

**Authors:** Rhys A. Farrer

## Abstract

Secretion signals are ancient and functionally conserved sequence motifs that orchestrate function and intended destination of cleaved encoded proteins (*1–3*). To investigate the genomic landscape of secreted proteins, 4,694 annotated eukaryotic genome assemblies were analysed. Genes encoding secretion signals (*n* = 5.2 million) were consistently enriched in genomic regions with longer flanking intergenic regions (FIRs). Consecutive genes with characteristic FIR lengths were enriched for genes with secretion signals. Intriguingly, many eukaryotic pathogens and parasites have the most significant association between genes encoding secretion signals and their intergenic distance. Almost every category of repeat was found in greater number flanking genes encoding secretion signals, with especially strong enrichment of simple, unknown, and low complexity repeats in fungal genomes. Despite higher repeat counts, the total repeat length was consistently shorter around genes with secretion signals, suggesting a prevalence of truncated or fragmented repeats in these regions. Several GO-terms assigned to genes with secretion signals were consistently enriched across genome assemblies in each kingdom. Common GO-enrichment patterns were also identified in genes categorised by their FIR. These results hint at an anciently conserved genomic architecture and mode of evolution in eukaryotes, characterised by long FIRs and fragmented repeat landscapes, likely driven by mechanisms such as repeat-driven gene copy number variation (*4*), differential mutation rates (*5*) and chromatin remodelling (*6*). This conserved association highlights the potential of genome structure to drive innovation in secreted protein function.

## Introduction

Secreted proteins are broadly conserved and functionally important across all domains of life. In eukaryotes, they are typically targeted to the extracellular space via N-terminal secretion signals and processed through the endomembrane system. In prokaryotes, a wide range of specialised secretion systems are used including Sec, Tat, and Type I-VII, facilitating protein export across the cell membrane. These systems, while mechanistically distinct from the eukaryotic endomembrane system, reflect the deep evolutionary origins of protein secretion and its essential role in cellular function and environmental interaction(*7, 8*).

Secreted proteins serve pivotal roles in the development of multicellular organisms, disease and infection, acting as structural matrix, extracellular enzymes, and signal molecules (*1*). The default pathway for eukaryotic protein secretion is the Golgi pathway, initiated by translocation of the protein through the endoplasmic reticulum (ER) (*2*). The process begins when free ribosomes translate mRNA destined for the secretory pathway, the first 5-30 amino acids (N-terminal) of which encode a signal peptide. The signal peptide is recognised and bound by a signal recognition particle, leading to elongation arrest and targeting the ribosome-nascent chain complex to a translocation channel in the ER membrane (*3*). Once inside the ER, the signal sequence is cleaved by a signal peptidase, folded and directed to its intended location, which is governed by differences in the amino acid characteristics of the signal peptide. Cleaved proteins from the secretory pathway can be directed to other organelles, secreted from the cell, or inserted into cell membranes or the cell wall.

Many components of the secretory pathway including signal peptides, signal recognition particles (SRPs), and the endoplasmic reticulum targeting machinery are functionally conserved across the super-kingdoms of life (*3, 9*), suggesting an ancient origin despite rapid sequence evolution of individual signal peptides. Indeed, secreted proteins are thought to evolve more rapidly than intracellular proteins, regardless of expression or interaction complexity, suggesting unique evolutionary pressures linked to subcellular localization(*10*). While expression level across all genes is inversely correlated with evolutionary rate (E-R anticorrelation), this correlation may be weaker in genes encoding secretion signals, potentially due to the secreted protein having a reduced risk of intracellular misfolding or misinteractions (*11–13*). Secreted proteins have also been shown to have signatures of rapid evolution outside their signal peptide in diverse taxa including both prokaryotes (*14*) and eukaryotes (*15, 16*). Together, these patterns highlight the evolutionary distinctiveness of secreted proteins and raise the question of whether this is reflected in their genomic positioning and broader genome architecture.

Secreted effector proteins that target host immune and recognition proteins, as well as pathogenicity factors encoded by several plant and animal pathogens have been recognised to associate with repeat-rich regions of their genome(*15, 16*). Furthermore, genes with secretion signals have characteristic longer flanking intergenic distances (*15, 16*), which are hypothesized to drive their evolution through copy number variations and diverse epigenetic and chromatin mechanisms such as differences in accessibility to repair mechanisms, homologous recombination, cytosine methylation, and repeat induced mutagenesis.

To determine if eukaryotic secreted proteins broadly associate with repeat-rich genomic regions, available high-quality and annotated eukaryotic genomes from NCBI GenBank were downloaded and screened for secretion signals, identifying 5.2 million genes with putative secretion signals. Animal genomes encoded the highest proportion of secreted proteins, while plants had the lowest proportion. Genes encoding secretion signals had significantly longer median 3’ and 5’ flanking intergenic regions (FIR, i.e., the non-coding genomic sequences immediately upstream and downstream of a gene) compared with those that didn’t. To visualise this, the 3’ and 5’ FIR of each gene was plotted (x-axis and y-axis, respectively), and classified into four FIR quadrants based on whether their 3′ and 5′ intergenic distances were above or below the median values of all genes: upper-right (Q_UR_), upper-left (Q_UL_), lower-right (Q_LR_), and lower-left (Q_LL_). Genes encoding secretion signals were highly enriched in the Q_UR_ quadrant, indicating consistently longer flanking intergenic distances. Furthermore, those genomic regions contained significantly more repetitive elements, especially in fungi, yet these repeats were consistently shorter in total length. Together, these results demonstrate an anciently conserved association with genes encoding secreted proteins and repetitive elements.

The correlation between gene function and FIR suggest that the genomic localisation of those genes is not incidental. Secreted proteins, such as those involved in host-pathogen or host-environment interactions, often face strong selection to diversify. This raises the possibility that their positioning within repeat-rich, sparsely populated genomic regions reflects an adaptive strategy to promote evolutionary flexibility, enabling regulatory independence, and gene family expansions.

## Results

All publicly available annotated eukaryotic genome assemblies from NCBI GenBank were downloaded and analysed, spanning the four major eukaryotic superkingdoms based on their entry in the NCBI Taxonomy Browser (Animal, Plant, Fungi and Other:). After removing low quality assemblies (**Figs. S1-S8**), a phylogenetic tree based on core eukaryotic genes was constructed to visualise the breadth of taxonomic sampling and the diversity of secreted protein abundance across the eukaryotic domain (**Fig. 1A; Dataset S1**). The tree highlights both the overrepresentation of fungal genomes in public databases and the broad variability in the proportion of secretion signals among species, including closely related taxa, reinforcing the complexity and ecological diversity of eukaryotic secretomes.

**Figure 1.**
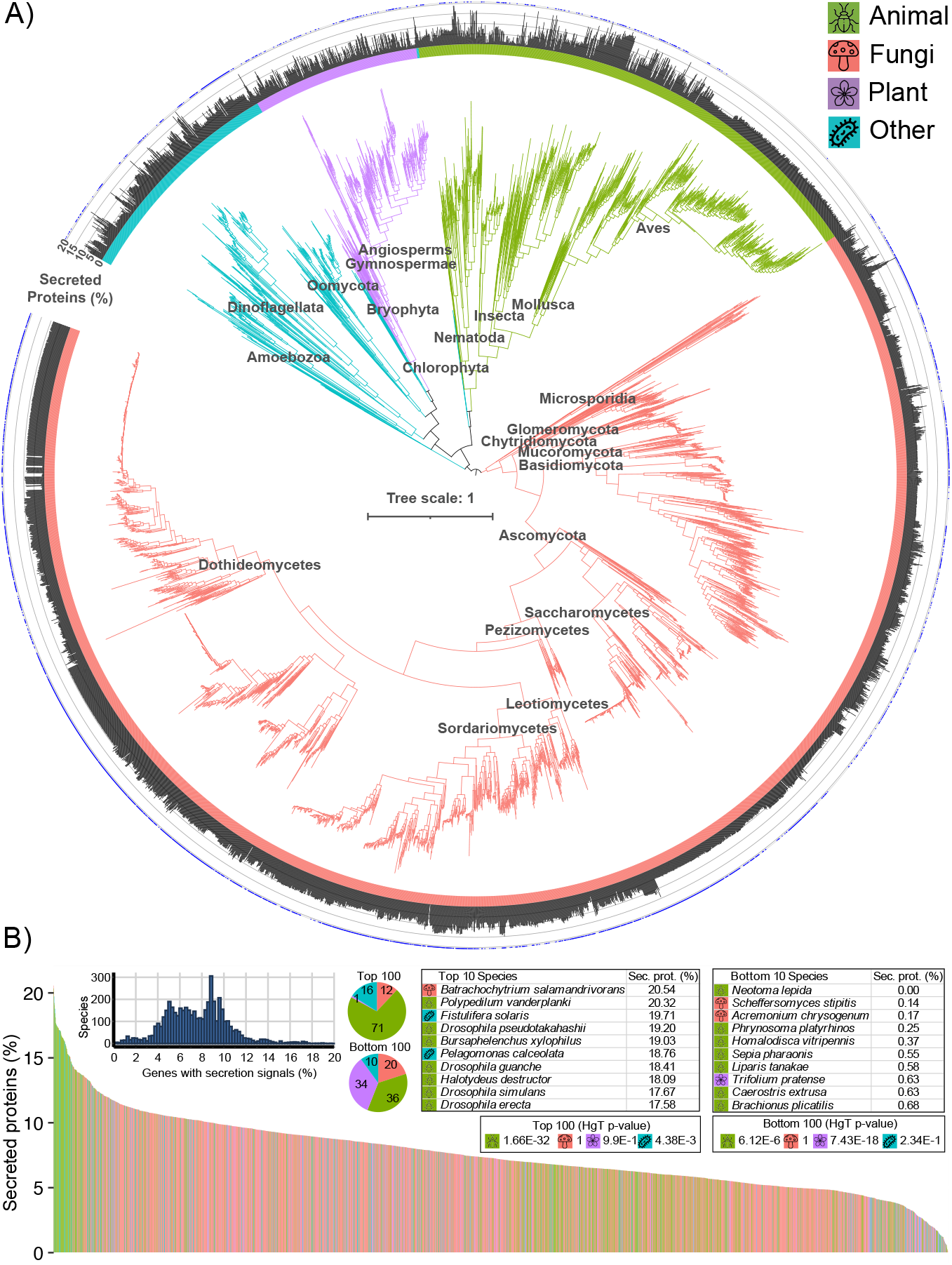
**A)** A midpoint rooted phylogenetic tree of 4,694 eukaryotic genome assemblies, constructed from multiple sequence alignments of 125 core eukaryotic genes (alignment length (amino acids and gaps) = 714,761 per taxa) and inferred using FastTree. Various sub-kingdom taxonomic groups have been added of major groups above the nodes for illustrative purposes. The inner circle shows the kingdom, followed by a barchart showing the percent of secreted proteins for each assembly. The outside circle shows a blue dot for each assembly that has a significant enrichment of genes with secretion signals in Q_UR_. **B)** Percentage of genes encoding secretion signals across 4,694 eukaryotic genome assemblies, ordered from highest to lowest. For species with multiple assemblies, panel B includes a single representative assembly per species selected as the assembly with the highest percent of predicted secreted proteins. The top left panel shows a histogram of species counts by percentage of genes with secretion signals. Pie charts display the kingdom composition for the top 100 and bottom 100 genomes based on secreted protein percentages. Tables list the top 10 and bottom 10 species. Results of 4 hypergeometric tests (upper tail *p*-value) assess kingdom-level overrepresentation in the top/bottom 100 sets (bold indicates significance). Green = animal, red = fungi, purple = plant, blue = other.

Between 0 and 21% (mean□=□7.55%, median□=□7.54%, σ□=□2.81) of genes per assembly encode predicted secretion signals. Overall, 7.1% of all eukaryotic genes analysed contain a predicted secretion signal (**Figs. 1B; S9; Dataset S2**). Durum wheat has the greatest number of genes with secreted signals (*n* = 18,752/190,470; 9.8%) followed by bread wheat (17,686/135,934; 13%). The amphibian pathogen *Batrachochytrium salamandrivorans* has the largest percent of genes with secretion signals (*n* = 2,353/11,454; 20.54%). Across the eukaryotes, only *Neotoma lepida* (the desert woodrat) was predicted to encode no proteins with secretion signals among its 24,320 annotated protein encoding genes. This result is likely due to annotation or prediction artifacts, rather than a true biological absence of secreted proteins. Two-tailed *t*-tests suggests significant differences between the percent of genes with secretion signals compared with genes without secretion signals across the animals, fungi, plant and other (*p* = 0.01, 3.04 × 10^-5^, 7.32 × 10^-18^ and 1.27 × 10^-5^, respectively). The discrepancy between percent of secreted proteins across taxonomic ranks and species underscores their diverse ecological roles.

Secreted and non-secreted proteins are encoded by genes of similar lengths, with near-identical medians (1,050 vs 1,056 bp, respectively) and slightly shorter mean lengths (1,267 vs 1,343 bp, respectively). Given the very large sample size (>58 million genes), even minor trends are likely to reach statistical significance. Accordingly, only weak yet significant correlations were observed between gene length and intergenic distances: 5′ upstream flanking intergenic regions (FIRs) (Pearson’s r = -0.0116, *p* < 2.2 × 10-^16^, 95% CI: -0.01188 to -0.01137) and 3′ downstream FIR (r = -0.0039, *p* < 2.2 × 10-^16^, 95% CI: -0.00367 to -0.00418). These small differences in gene length and flanking distance suggest that gene length is not a confounding variable in subsequent analyses and is unlikely to reflect meaningful biological differences on its own.

To address the potential link between genome architecture and intergenic spacing, the proportion of the genome annotated as coding sequence (coding fraction) and gene density (genes per megabase) for each of the 4,694 genome assemblies were calculated. Both metrics were negatively correlated with the length of 5′ and 3′ median flanking intergenic regions (FIRs), suggesting that genomes with higher gene densities or more compact gene organisation tend to have shorter intergenic regions. Specifically, coding fraction was negatively correlated with 5′ and 3’ FIR (r = -0.549 and -0.552, respectively; both *p* < 2.2×10^−1^□). Similarly, gene density showed significant negative correlations with both 5′ FIR and 3′ FIR (r = - 0.554 and r = -0.559, respectively; both *p* < 2.2×10^− 1^□). These findings confirm that, as expected, genome compaction is inversely related to FIR length.

Genes with secretion signals had longer FIRs than those without, with a 321 nt increase in 3′ median FIR (1,534 nt vs. 1,213 nt) and a 31 nt increase in 5′ median FIR (1,025 nt vs. 994 nt) (**Fig. 2A-B**). Across all eukaryotic genes, those encoding secretion signals (*n* = 5,213,813) were highly enriched in regions with above median 3′ and 5′ FIRs compared to all other genes (*n* = 68,128,352; hypergeometric test *p* < 2.22 × 10^-308^) (**Figs. 2C; S10**). A similarly strong enrichment was also observed in Quadrant Lower-Right (Q_LR_). The majority (58.4%) of eukaryotic genome assemblies had more genes with secretion signals in Q_UR_ than Q_LL_, and 2,483 (53%) showed a significant enrichment in Q_UR_ (*p* < 2.13 × 10^-6^) (**Fig. 1A**). Q_UR_ enrichment for genes with secretion signals was widely present across the four kingdoms, with a notable exception of the Aves, which only had 3 assemblies with this enrichment. Bootstrap analysis of randomly sampled genes supported that this level of enrichment was not due to chance (**Fig. 2D**). Notably, no correlation was found between the proportion of genes with secretion signals in an assembly and their enrichment among any of the quadrants (**Fig. S11**), suggesting that FIR-based enrichment is an independent genomic feature rather than a byproduct of secretion signal abundance. Together, these findings reveal a new general association between genes encoding secretion signals and longer FIR.

**Figure 2.**
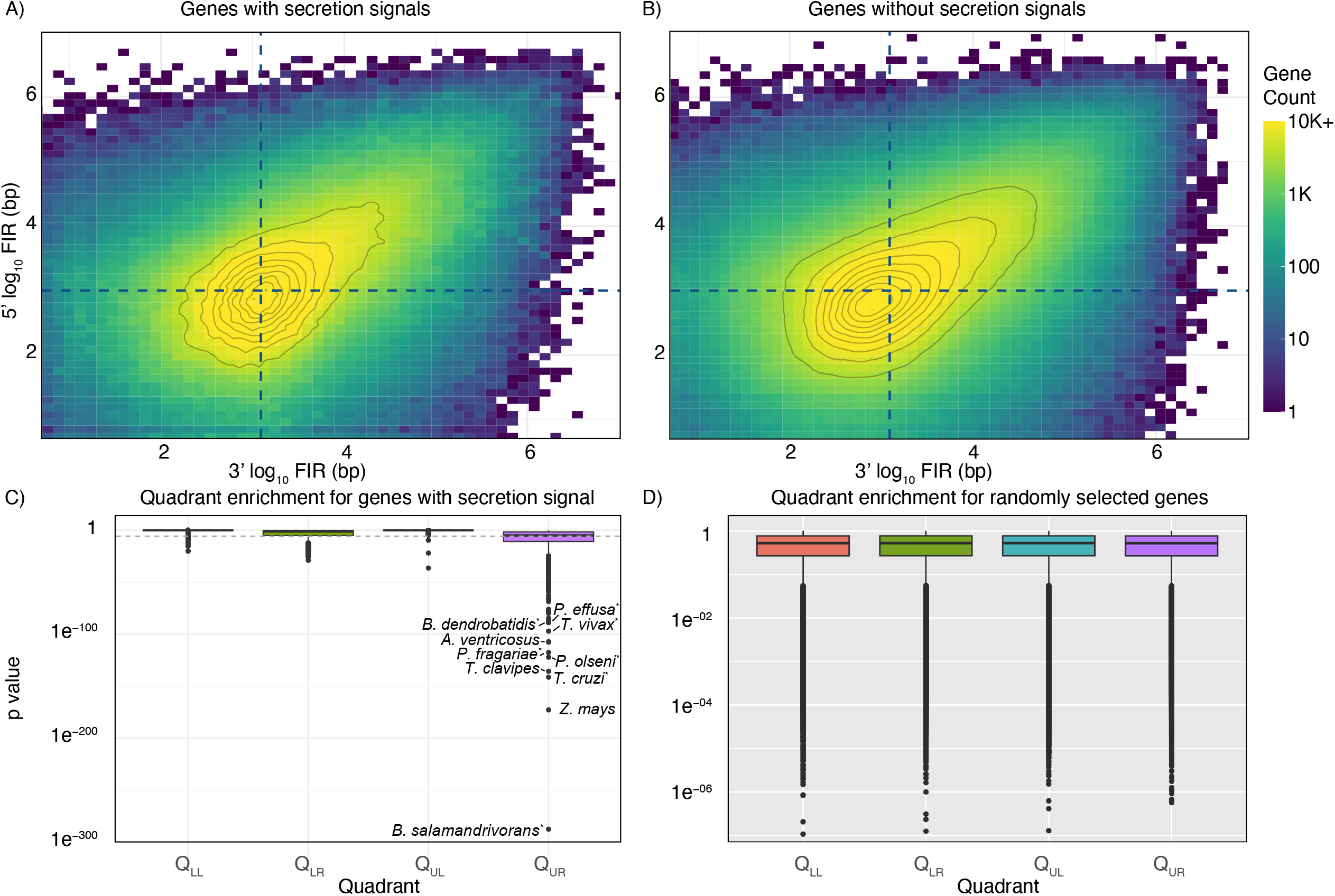
(**A-B**) 3’ and 5’ flanking intergenic distances (FIR) distances for eukaryotic genes. Dotted lines indicate the median FIR values for all genes, which define four quadrants: upper left (UL), upper right (UR), lower left (LL) and lower right (LR). **A**) Genes with secretion signals. **B**) Genes without secretion signals (downsampled to 4,372,912 random genes, to normalise to the number of genes with secretion signals). **C**) Enrichment of genes with secretion signals in each FIR quadrant across 4,694 genome assemblies. Upper-tail hypergeometric *p*-values are plotted for each genome. The ten most significant enrichments (all in the upper right quadrant, UR) are labelled; seven of these correspond to known pathogens (indicated by asterisks). **D**) Control analysis showing hypergeometric test *p*-values for randomly selected genes across genome assemblies (1,000 bootstraps per assembly), validating enrichment patterns observed in panel C.

To evaluate whether consecutive genes (regardless of whether they encode secretion signals or not) tend to occupy the same FIR-based quadrant more often than expected by chance, I applied a custom time-discrete Markov model, accounting for the number of genes in each quadrant on each contig. Using a Bonferroni-adjusted significance threshold (*p* < 1.97 × 10^-10^), I identified 615 significant clusters: 284 in Q_UR_ (long 5′ and 3′ FIRs) and 331 in Q_LL_ (short 5′ and 3′ FIRs), each comprising 14 to 217 consecutive genes (**Dataset S3**). The top three most significant Q_UR_ gene clusters were all found in pathogens: 45 consecutive Q_UR_ genes on Chromosome 18 of *Leishmania major* Friedlin (*p* = 9.86 × 10^-31^), 59 on Chromosome 13 of *Plasmodium malariae* (*p* = 5.20 × 10^-27^), and 50 on Chromosome 14 of *Plasmodium yoelii* (*p* = 7.41 × 10^-25^). Notably, *Plasmodium* species contributed 48 of the 284 Q_UR_ clusters (17%) but showed no enrichment for Q_LL_ clusters. Genes within Q_UR_ clusters were strikingly enriched for secretion signals (38,994 of 447,407; 8.7% encoded secretion signals; hypergeometric test, *p* < 2.22 × 10^-308^) whereas Q_LL_ clusters showed no enrichment (25,162 of 449,136; 5.6%; *p* = 1). Together, these results support the existence of genomic sub-regions characterised by expanded intergenic spacing that are significantly enriched for genes encoding secretion signals, particularly in pathogenic lineages.

To investigate the composition of the FIR among eukaryotic genomes, repeats were predicted for a subset (8.4%) of all genome assemblies (**Table S1; Datasets S4-S5**). Genome length and N50 (as a metric for contiguity/quality) showed weak-to-moderate but statistically significant correlations with total repeat content (%) (Genome length vs repeat %: Pearson r = 0.371, R^2^ = 0.138, *p* = 2.81E^-14^, 95% CI [0.283, 0.453], N50 vs repeat %: Pearson r = 0.392, R^2^ = 0.153, *p* = 7.31E^-16^, 95% CI [0.305, 0.472]). However, the association was stronger among individual kingdoms, with genome length correlating with repeat content (%) as much as R^2^ = 0.63 for plants (**Fig. S12**). Consistently across each superkingdom, repeat elements were significantly more numerous when flanking genes with secretion signals, with the strongest enrichments observed in fungi (**Fig. 3**). *t*-tests revealed that simple repeats, low-complexity elements, and repeats of unknown classification were particularly enriched near genes with secretion signals in fungal genomes (adjusted *p* < 0.001) (**Tables S2-3**). Although genes encoding secretion signals are flanked by a greater number of repeat elements, the individual repeats surrounding genes lacking secretion signals are consistently longer, a pattern found for both repeats upstream and downstream of genes with or without secretion signals, with and without normalisation for the number of those genes (**Figs S13-S17, Tables S4-S6**). Together, this reveals that the repeat landscape around genes with secretion signals is enriched yet fragmented, dominated by shorter, possibly truncated or degenerate elements, while repeats near genes lacking secretion signals are fewer but significantly longer.

**Figure 3.**
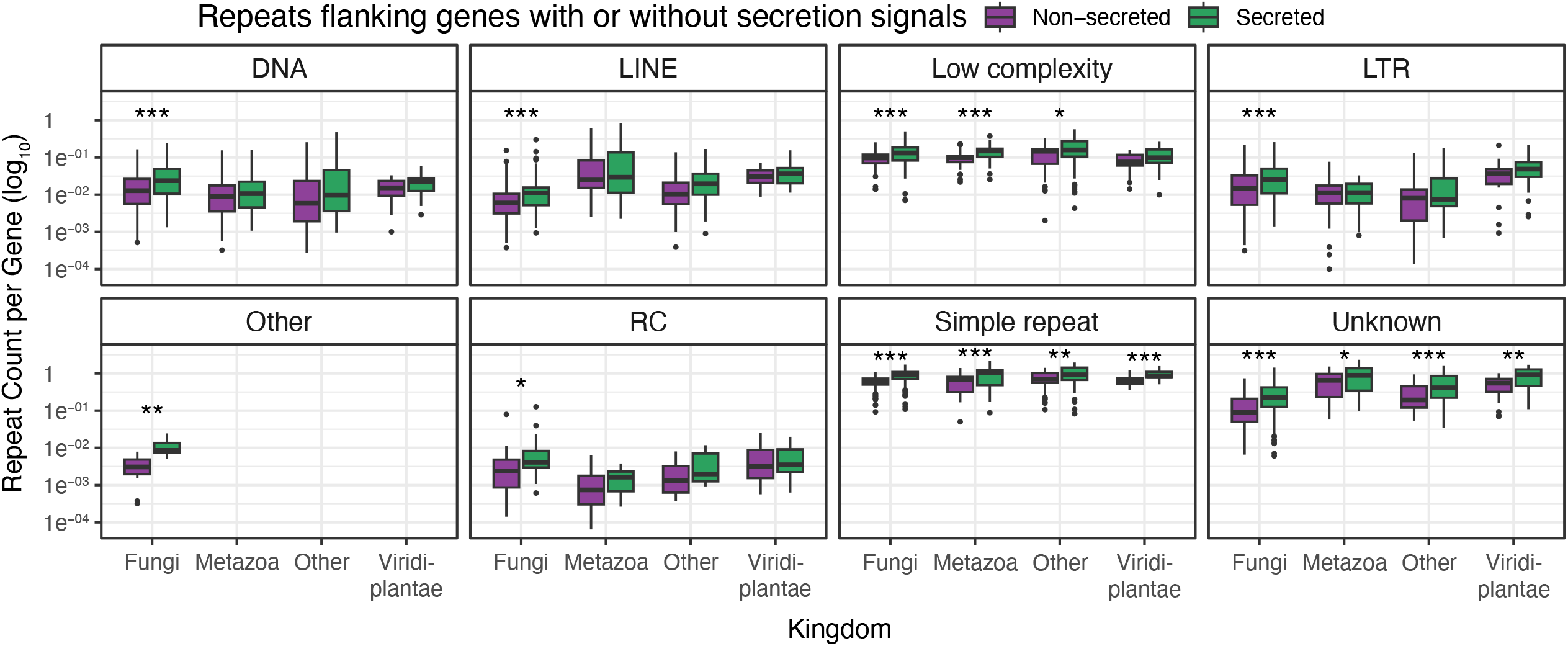
The abundance of repetitive elements flanking genes with (green) or without (red) secretion signals across different repeat superclasses categorised by RepeatModeller. “Other” repeats are repeat elements that are known, but RepeatModeller has not been able to confidently assign to one of the major classes. Repeat counts were normalised by the number of genes in each category (secreted vs non-secreted) per genome assembly. Asterisks indicate statistical significance based on *t*-tests, corrected for multiple testing using the Benjamini-Hochberg (BH) method. *** = *p* < 0.001, ** = *p* < 0.01, * = *p* < 0.05.

A pattern of significant functional enrichment of GO terms (defined as q-value < 0.01) was identified in genes encoding secretion signals vs genes lacking secretion signals. Among genes with secretion signals, approximately 87% of fungal genomes and 74% of animal genomes had significant enrichment of GO:0005524 (ATP binding), 87% of plant genomes had significant enrichment of GO:0099503 (secretory vesicle), and 45% of genomes in the ‘other’ kingdom category had significant enrichment of GO:0005788 (endoplasmic reticulum lumen) (**Figure 4; Dataset S6**). From each kingdom, the most significantly enriched GO term (ordered by *q*-value) among secreted vs non-secreted proteins in any genome assembly was selected. I then plotted the FIR for all proteins that had that GO term assigned to investigate correlation between GO terms enriched in secreted proteins and their FIR (**Datasets S7, S8**).

**Figure 4.**
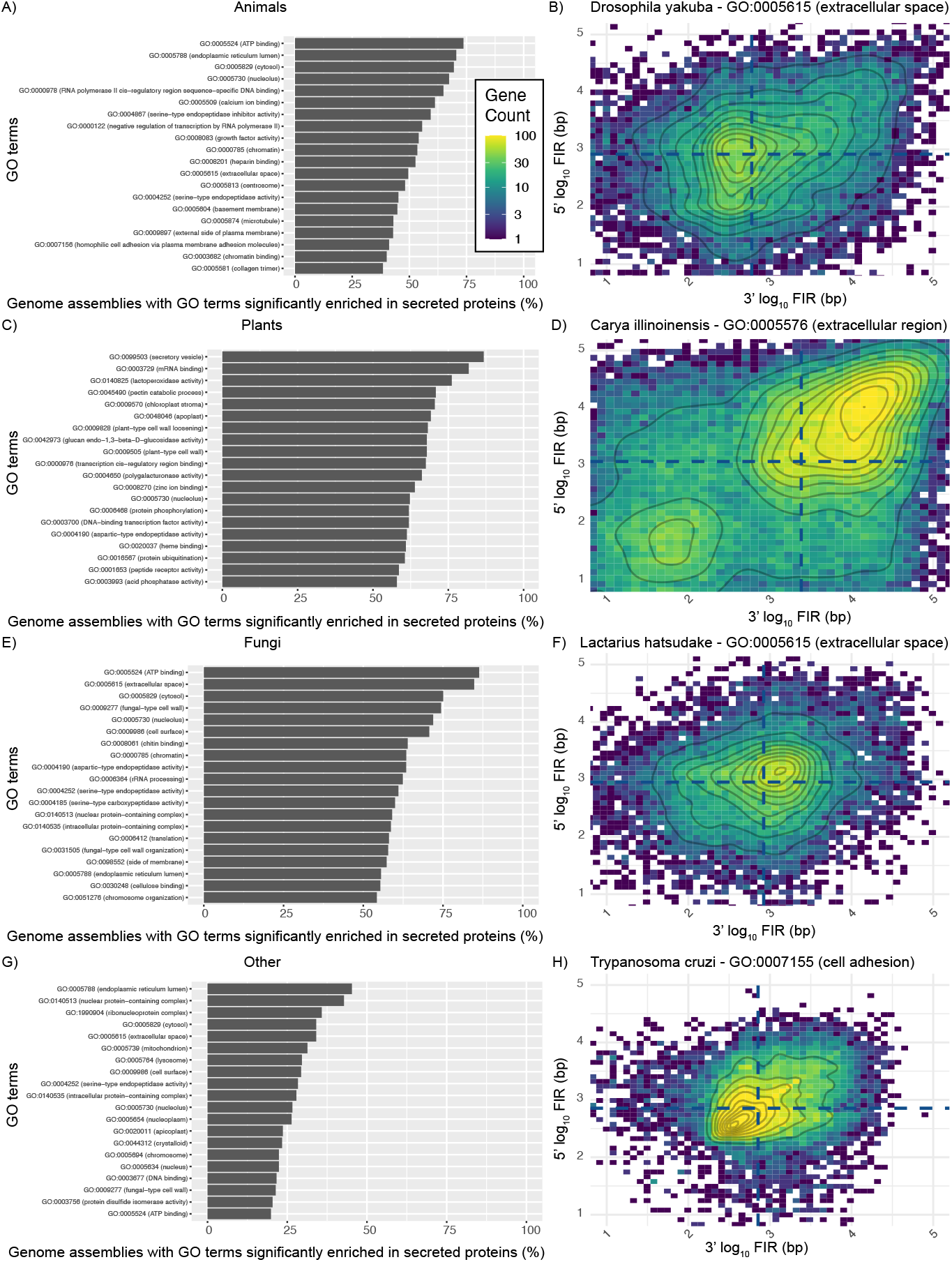
GO-terms enriched among secreted proteins compared with non-secreted proteins. (**A, C, E, G**) For each genome assembly in each kingdom, GO enrichment was tested for secreted versus non-secreted proteins. Bars show the 20 GO terms that most frequently reached significance across assemblies in that kingdom. The x-axis indicates the percent of assemblies in which each GO term was significantly enriched in secreted proteins. (**B, D, F, H**) Illustrative examples. For each kingdom, the single assembly and GO term combination with the lowest adjusted p-value (FDR) was selected, and the 5′ and 3′ flanking intergenic region (FIR) lengths are plotted for all genes annotated with that GO term in the selected assembly (assembly and GO term shown in each panel title). Heatmap colour denotes gene counts. Dashed lines indicate the median 5′ and 3′ FIR lengths for all genes in that assembly, showing that GO term associated genes can be shifted relative to the genome-wide median.

Among genes with secretion signals, the most significantly enriched GO-term in animals (*Drosophila Yakuba*; GO:0005524; ATP binding; *q* = 7.75E-152), was not significantly enriched in any of the four quadrants (**Figures 4B**). Among genes with secretion signals, the most significantly enriched GO-term in plants (*Carya illinoinensis*; GO:0005576; extracellular region; *q* = 6.47E-185) was also significantly enriched in Q_UR_ (*q* = 3.96E-20; **Figures 4D**). Indeed, of the 46 terms enriched in Q_UR_ for C. *illinoinensis*, ‘extracellular region’ was the 7^th^ most significantly enriched, behind ‘heme binding’, ‘iron ion binding’, ‘oxidoreductase activity’, ‘DNA-binding transcription factor activity’, ‘response to inorganic substance’, and ‘defence response to fungus’. Among genes with secretion signals, the most significantly enriched GO-term in fungi (*Lactarius hatsudake*; GO:0005615; extracellular space; *q* = 1.20E-114) was also significantly enriched in Q_UR_ (*q* = 3.51E-10; **Figures 4F**), and of the only 7 terms identified, was 2^nd^ most significantly enriched after tripeptidyl-peptidase activity (*q* = 7.03E-11). Lastly, among genes with secretion signals, the most significantly enriched GO-term in the ‘other’ kingdom category (*Trypanosoma cruzi*; GO:0007155; cell adhesion; *q* = 9.91E-126) was not significantly in any of the four quadrants, despite appearing enriched in the Q_LL_ (**Figures 4H**). Therefore, some GO term enrichments among secreted proteins are also found when examining only QUR (and not secreted profile).

Quadrant enrichment for GO-terms irrespective of encoding secretion signals was also investigated (**Datasets S8 and S9**). The most frequently enriched GO-terms in the Q_UR_ were ‘DNA-binding transcription factor activity’ in animals (64%), ‘DNA-binding transcription factor activity’ in plants (77%), ‘plasma membrane’ in fungi (21%) and ‘nucleus’ in ‘other’ (11%) (**Figure S18**). The most significant terms in Q_UR_ for assemblies belonging to each of the four kingdoms was *Trichonephila clavipes* GO:0005515 (protein binding), *Mucuna pruriens* GO:0005515 (protein binding), *Rhizophagus irregularis* DAOM 181602 GO:0006468 (protein phosphorylation), and *Plasmodium sp. gorilla* clade G1 GO:0020033 (antigenic variation) (**Figure S18**). Together, these reveal common patterns of function across eukaryotic genes encoding secretion signals and genes categorised based on their intergenic distance.

## Discussion

Intergenic distances and gene function have been previously associated with genome organisation in “two-speed genomes”, which is a concept describing genomes partitioned into gene-dense, conserved regions that are more conserved (or have a slower evolutionary speed) alongside other genomic regions that are gene-sparse and repeat-rich that often include rapidly (or faster) evolving genes. This genomic feature has been described in pathogens such as *Batrachochytrium salamandrivorans* (*15*) and *Phytophthora infestans* (*16*). Here, by analysing 4,694 annotated eukaryotic genome assemblies, I demonstrate that genes encoding secreted proteins consistently reside in genomic regions with longer flanking intergenic regions (FIRs). This pattern appears to be a general feature across eukaryotes. Because public databases contain uneven numbers of assemblies per species, the results were mostly interpreted as broad trends across QC-passing assemblies. Focusing on species-collapsed sensitivity analyses represent a useful direction for future work. Localisations of genes encoding secretion signals may be the result of selective pressures favouring gene family expansions, changes in chromatin accessibility or regulatory changes, driven by mechanisms such as repeat-driven gene copy number variation (*4*), differential mutation rates (*5*) and chromatin remodelling (*6*). Understanding how individual gene families contribute to these patterns would likely provide additional insights, such as by highlighting a particular gene function or gene family driving that pattern.

Among eukaryotes, animals have the overall highest proportion of genes encoding secretion signals. Nevertheless, notable non-animal outliers reinforce the biological relevance of these patterns across diverse kingdoms and lineages. For example, the amphibian pathogen *Batrachochytrium salamandrivorans* encodes the highest proportion of secreted proteins and has a significant enrichment of those secreted proteins in the upper-right (Q_UR_) FIR quadrant. Similarly, *Trypanosoma cruzi, Perkinsus olseni*, and *Phytophthora fragariae*, which are pathogens known for secreting host-interacting proteins, also showed strong Q_UR_ enrichment for genes with secretion signals. These species encode effectors and secreted enzymes that directly manipulate host environments, and their enrichment in high-FIR regions suggests that FIR structure may underlie or facilitate adaptive secretion strategies (*17–20*).

Strikingly, 53% of all assemblies surveyed exhibited Q_UR_ enrichment for genes with secretion signals, a pattern present across each major eukaryotic kingdom. Aves were a notable exception, showing an almost complete absence of this enrichment: among 339 avian genomes, only three assemblies showed Q_UR_ enrichment for genes with secretion signals: the common pigeon (*Columba livia*), the society finch (*Lonchura striata domestica*), and the Picui ground dove (*Columbina picui*). The otherwise widespread association of genes with secretion signals and longer FIR points to a previously underappreciated conserved genomic architecture for secretory function, wherein secretion-associated genes preferentially localise to genomic regions marked by longer flanking intergenic distances with more numerous repeat elements, potentially thereby impacting regulation and gene expression. The rapid evolution of secreted proteins previously observed to be faster than that of intracellular proteins across multiple taxa(*10*) may also be facilitated by their localisation within genomic regions enriched for repeats and long intergenic distances.

Gene clusters comprising consecutive genes belonging to a given quadrant had a significant enrichment of secretion signals. These gene clusters were widespread across the Eukaryota, but particularly prominent in pathogens. For example, *Leishmania* and *Plasmodium* species harboured the most statistically significant clusters, with *Plasmodium* species alone accounting for 17% of all Q_UR_ gene clusters identified across the Eukaryota, yet accounting for no significant Q_LL_ gene clusters, revealing directional bias in genomic organisation. These co-localised clusters of genes with secretion signals with extended intergenic spacing further suggest a non-random genome architecture characteristic for a range of primarily pathogenic species, as has been described for effector families in plant-pathogenic fungi including *Phytophthora infestans* (*16*), *Blumeria graminis* (*21*), *Zymoseptoria sp*. (*22*) and *Leptosphaeria maculans* (*23*). To detect these clusters, I implemented a discrete-time Markov chain model that computes the probability of observing uninterrupted runs of genes in each quadrant, accounting for their background frequency. While this is distinct from autocorrelation analysis, which measures correlation over fixed lags (i.e., whether values at one position in a sequence are correlated with values a fixed number of steps away, such as between adjacent or every n^th^ gene), the Markov approach was chosen to sensitively detect localised enrichment of consecutive genes.

Consistent with these patterns of longer FIRs around genes with secretion signals, those genes were significantly more likely to be flanked by repeat elements. Fungi in particular exhibited strong enrichment for simple, low-complexity, and unclassified repeats around genes with secretion signals. Although genes with secretion signals flanked a greater number of repeats, the individual repeats were consistently shorter compared to those flanking genes lacking secretion signals. This suggests a fragmented repeat landscape surrounding genes with secretion signals, possibly reflecting truncation, decay, or TE-associated rearrangements in these regions. For example, in *Batrachochytrium salamandrivorans*, large families of secreted metalloproteases are enriched for flanking LINE elements that have lost their functional domains (*24*). Such fragmentation may reflect transposon activity followed by subsequent selection to remove those elements.

The consistent localisation of secretion-associated genes to repeat-rich, sparsely populated genomic regions may reflect an adaptive strategy to promote rapid evolutionary change. Genes involved in host interactions, such as effectors that interact with the host immune system, tend to evolve quickly in response to host immune pressures or environmental variability. Genes in “higher-evolutionary-speed” regions (characterised by dynamic, repeat-rich, and less constrained architecture) may facilitate such adaptability by impacting mutation rate or type, promoting recombination, and allowing for gene copy number variation. Additionally, the extended intergenic distances may enable finer regulatory control by isolating transcriptional responses and reducing interference from neighbouring genes. This modular architecture could be especially advantageous in pathogens and symbionts, where flexible gene expression and rapid innovation are critical for survival.

Together, these findings provide evidence for a genomic niche for genes encoding secreted signals, characterised by long FIRs, repeat-rich yet fragmented or shorter repeat families, and frequent clustering with other secretion-encoding genes. This organisation may facilitate regulatory independence of individual genes in those clusters, enable rapid lineage-specific gene expansion, and allow adaptive diversification, especially in pathogens or symbionts interacting with complex host ranges or new hosts. Repeat-driven genome plasticity has been previously implicated in the expansion of gene families such as RXLR effectors in *Phytophthora* (*16*), HASP/SHERP genes in *Leishmania* (*25, 26*), and fungal LysM effectors (*27*).

Here, the broader evolutionary importance of repeat-mediated dynamics on genes with secretion signals is investigated, suggesting a common genomic architecture that facilitates functional innovation.

## Methods

Full details are given in SI Appendix. Briefly, all genome datasets were downloaded from NCBI GenBank on the 18^th^ of April 2022. Taxonomy ID was used to identify and split every file by their superkingdom (based on their entry in the NCBI Taxonomy Browser (*28*)). Duplicate gene entries were excluded from the annotation. Assemblies with < 182 annotated genes were excluded from the analysis. Core Eukaryotic Genes (CEGs) (*29*) coverage was used to further assess quality of assemblies and annotation (**Figures S1-S6**). 79 eukaryotic genomes were excluded because fewer than 50% of their protein encoding genes with ≥ 70% alignment coverage against the CEGs, indicating poor annotation or incomplete assemblies. After quality control, 4,694 eukaryotic genome assemblies remained for further analysis, with a combined 73,334,690 protein coding genes. Genes encoding secretion signals were predicted using SignalP 4.1 (*30*). All code for this project has been made available from https://github.com/rhysf/2speed_genomes, including calculating upstream and downstream distances and hypergeometric tests for enrichment. Repeat elements were predicted in 393 genome assemblies with RepeatMasker version 4.1.2-p1 (*31*) using *de novo* repeat libraries generated from each genome assembly with RepeatModeler version 2.0.3 (*32*) with LTRStruct.

For phylogenetic analysis, 125 core eukaryotic genes (CEGs) (*29*) were identified in 4,694 genome assemblies using blastp v2.2.24 (*33*) (parameters -evalue 1e-10 -max_hsps 5). Multiple sequence alignments were made using MUSCLE v3.8.31 (*34*) and refined with trimAl v1.4.1 (*35*). A phylogenetic tree was constructed with FastTree v2.1.11 (*36*). The tree was visualised using iTol v7 (*37*) with midpoint rooting. A Markov chain was used to calculate probabilities of consecutive numbers of genes in each Quadrant (as described above, each quadrant is defined by the median 3’ and 5’ FIR of all genes). Hypergeometric tests were used to evaluate enrichment of genes with secretion signals among each quadrant. To test the validity of the results, the number of genes encoding secretion signals was used to randomly select genes across each eukaryotic genome assembly, and their intergenic distances compared with the remaining genes using the same hypergeometric test approach. The process was repeated 1,000 times per genome. To assess whether certain repeat families were enriched or depleted flanking genes with or without predicted secretion signals, repeat counts were compared using *t*-tests and Wilcoxon rank-sum tests with Benjamini-Hochberg (BH) correction. Functional predictions were assigned to all protein sequences using Diamond2GO(*38*), using the V1 database, and enrichment determined by Wilcoxon rank-sum tests with BH correction.

## Acknowledgements

Thanks to useful discussions with Theresa Wacker, Hugh Gifford, Nicholas Helmstetter and David Studholme. Thanks to Ben (user 173082 of stats.stackexchange.com) for description and solution to implementing the Markov chain algorithm (question 26988).

