## Supplemental Information for "Eukaryotic secreted proteins are encoded in repeat-rich genomic regions"

Rhys A. Farrer^1,*^

^1^Medical Research Council Centre for Medical Mycology at the University of Exeter, Exeter, United Kingdom

**Table of contents**

**Supplementary Methods … 2**

**Supplementary Results … 6**

**Supplementary Figures … 11**

**Supplementary Tables … 23**

**Supplementary Dataset Legends … 25**

**Supplementary References … 26**

**Supplementary Methods**

**Genomic data curation and QC**

All genome datasets (FASTA and GFF3, *n* = 1,271,677 for each) were downloaded from NCBI GenBank based on the assembly_summary_genbank.txt (ftp://ftp.ncbi.nlm.nih.gov/genomes/ASSEMBLY_REPORTS/) starting on the 18^th^ of April 2022 and completing 2-3 weeks later. The Taxonomy ID from the assembly_summary_genbank.txt file was used to identify and split every file by their Superkingdom (Archaea, Bacteria, Eukaryota, Viruses, Unknown_superkingdom) based on their entry in the NCBI Taxonomy Browser ^1^. Genomes with < 182 annotated genes were excluded (11 Archaea, 316 Bacteria, 138 Eukaryota, 42,061 Viruses, 20 Other) based on investigation of non-viral genomes, and the fewest numbers of genes found in any prokaryote: *Carsonella ruddii* ^2^. Several species (*n* = 205) had duplicate entries (the same feature and description): these duplicate entries were excluded from those GFF3s. Only the 4,694 eukaryotic nuclear genomes were analysed.

To avoid redundancy due to multiple isoforms or genes with alternative splice variants, CDS annotations were collapsed into single loci by summing all exon/CDS coordinates per gene. Genes were then sorted and filtered for overlap, with any subsequent overlapping entries excluded. This retained only a single representative gene per locus, reducing redundancy while preserving the primary genomic context for intergenic distance and repeat content analyses.

Gene sequences were extracted using the descriptions of the CDS feature in the GFF3. Eleven GFF3 did not specify CDS, so the ‘gene’ annotation was used instead. Gene sequences were translated to protein sequences using the standard code (based on being applicable to most species). Quality Control (QC) was performed based on Blast+ blastp coverage of Core Eukaryotic Genes (CEGs) ^3^ (**Figures S1-S6**). 79 eukaryotic genomes were excluded based on having <50% of their protein encoding genes with ≥70% CEG coverage. One further genome assembly (*Trypanosoma rangeli* SC58 (GCA_000492115.1) was excluded from flanking intergenic distance analysis, owing to it having a highly fragmented genome (*N_contigs_* = 9,066; *N_Max_* = 25 Kb, *N_50_* = 2.1 Kb) resulting in only 7 genes with upstream and downstream, which included no genes with secretion signals. After QC, 4,694 eukaryotic genome assemblies remained for further analysis, with a combined 73,334,690 protein coding genes.

Genes encoding secretion signals were predicted using SignalP 4.1 ^4^, identifying 5,213,813 genes with secretion signals (**Dataset S2**). Code available from <https://github.com/rhysf/2speed_genomes> was used to calculate upstream and downstream distances of all genes that had a gene upstream and downstream (i.e., not at the ends of contigs, scaffolds or chromosomes). The intergenic distances for 4,432,286 genes with secretion signals out of a total 59,473,322 protein coding genes were calculated and compared. The same Github repository includes R code to perform hypergeometric tests for enrichment in the 4 quadrants defined by the flanking intergenic regions of subsets of genes.

Repeat calling was computationally demanding, preventing analysis of all 4,694 eukaryotic nuclear genomes, some of which required months of computational processing. However, the first 393/4,694 (ordered alphanumerically by GCA number) (8.4%) of genome assemblies were repeat called including 227 Fungi (7.2%), 55 Metazoa (6.4%), 29 Plants (9.6%), and 82 Other (20.4%) (**Datasets S4-S5**). A further 7 mostly small or fragmented genome assemblies (5 Other, 2 Fungi; *Cryptosporidium muris* RN66, *Cryptosporidium parvum* Iowa II, *Encephalitozoon intestinalis* ATCC 50506, Encephalitozoon romaleae SJ-2008, *Metarhizium robertsii* ARSEF 23, *Monosiga brevicollis* MX1, *Naegleria gruberi*) did not yield any repeats. Of those assemblies with no predicted repeats, the total genome assembly lengths excluding ambiguous bases ranged from 2.1 Mb - 39 Mb. Repeat elements were predicted in each genome assembly with RepeatMasker version 4.1.2-p1 ^5^ using *de novo* repeat libraries generated from each genome assembly with RepeatModeler version 2.0.3 ^6^ with LTRStruct.

Functional predictions were assigned to all protein sequences using Diamond2GO^7^, using the V1 database (nr_clean_d2go_20230720.faa.dmnd). Functional enrichments using Hypergeometric tests among genes encoding secretion signals compared with genes not encoding secretion signals were performed using R phyper (**Datasets S6 and S7**) where q-values < 0.01 were recorded. Functional enrichments of each GO-term among the four quadrants defined by FIR was also performed (**Datasets S8 and Datasets S9**) where q-values < 0.00001 were recorded and the generic GO:0003674 (molecular_function) was excluded from the results.

**Phylogenetics**

For phylogenetic analysis, 125 core eukaryotic genes (CEGs) ^3^ were identified from 4,694 genome assemblies (using the peptide sequences based on annotation files) with blast v2.2.24 ^8^ (parameters -p blastp -e 1E-10 -z 20000 -f 11 -F F -b 1 -v 4). Each CEG (comprising 4,694 individual sequences) were placed in separate files, and multiple sequence alignments made using MUSCLE v3.8.31 ^9^. Phylogenetically uninformative sites from the multiple sequence alignments were removed using trimAl v1.4.1 ^10^ with default settings. Each of the CEGs were concatenated together (leaving a single sequence for each genome assembly) and a phylogenetic tree constructed with FastTree v2.1.11 ^11^. The phylogenetic tree was visualised using iTol v7 ^12^ with midpoint rooting.

**Markov chain analysis and bootstrap analysis**

A Markov chain was used to calculate probabilities of consecutive numbers of genes characterised by a given quadrant based on its median 3’ and 5’ FIR (**Dataset S4**). Across each chromosome or contig, every gene was characterised as one of 4 outcomes (Q_UL_, Q_UR_, Q_LR_, Q_LL_). A sequence (*n*) of any given length (*k*) is considered exchangeable (although Q_LL_ cannot be flanked by Q_UR_). For any given value of *𝑛*, let *𝒲* be the event that the substring of interest occurs, and let *ℋ_𝑎_* be the event that the last *𝑎* outcomes are the first *𝑎*<*𝑘* characters in the substring of interest (but no more than this). We use these events to give the following partition of *𝑘*+1 possible states of interest:

$$\begin{matrix} State 0 & \bar{\mathcal{W}}\cap\mathcal{H}_{0}, \\ State 1 & \bar{\mathcal{W}}\cap\mathcal{H}_{1}, \\ State 2 & \bar{\mathcal{W}}\cap\mathcal{H}_{2}, \\ State 3 & \bar{\mathcal{W}}\cap\mathcal{H}_{3}, \\ \vdots& \vdots\\ State k-1 & \bar{\mathcal{W}}\cap\mathcal{H}_{k-1,} \\ State k & \mathcal{W} \end{matrix}$$

Since the sequence of outcomes is assumed to be exchangeable, we have independent outcomes conditional on their respective probabilities (𝜃Q_UL_ + 𝜃Q_UR_ + 𝜃Q_LR_ + 𝜃Q_LL_ = 1). This can be represented as a discrete-time Markov chain that begins in State 0 at *𝑛* = 0 and transitions according to a probability matrix that depends on the substring of interest. The transition matrix is a (*𝑘* + 1)(*𝑘* + 1) matrix representing the probabilities of transition using the above states. For example, if the substring is 2, 3 or 4 consecutive genes characterised as Q_UR_ (described below as Q for brevity), the transition matrices would be:

$$p=\left[ \begin{matrix} 1-\theta_{Q} & \theta_{Q} & 0 \\ 1-\theta_{Q} & 0 & \theta_{Q} \\ 0 & 0 & 1 \end{matrix} \right] p=\left[ \begin{matrix} 1-\theta_{Q} & \theta_{Q} & 0 & 0 \\ 1-\theta_{Q} & 0 & \theta_{Q} & 0 \\ 1-\theta_{Q} & 0 & 0 & \theta_{Q} \\ 0 & 0 & 0 & 1 \end{matrix} \right] p=\left[ \begin{matrix} 1-\theta_{Q} & \theta_{Q} & 0 & 0 & 0 \\ 1-\theta_{Q} & 0 & \theta_{Q} & 0 & 0 \\ 1-\theta_{Q} & 0 & 0 & \theta_{Q} & 0 \\ 1-\theta_{Q} & 0 & 0 & 0 & \theta_{Q} \\ 0 & 0 & 0 & 0 & 1 \end{matrix} \right]$$

Once the transition matrix is constructed, for a given value of *𝑛* the probability of having the substring in the chain is $P(W|n)=\{Pn\}0,k$. This probability is zero for all *𝑛* < *𝑘*.

While autocorrelation is a well-established method to quantify similarity between values a fixed number of steps away (lag), across a sequence, the aim was not to measure lag-dependent correlation but rather to detect significant runs of consecutive genes falling into the same FIR quadrant, particularly Q_UR_. The approach described above and implemented as a Perl script on the accompanying GitHub repo (https://github.com/rhysf/2speed_genomes/blob/main/perl_scripts/Markov_chain_from_quadrant_file.pl) modelled quadrant transitions along contigs as a discrete-time Markov chain. This allowed the exact probability of observing uninterrupted runs of specific quadrant types to be calculated under the assumption of independence, given the observed frequencies within each contig. This approach effectively identifies enriched gene clusters while accounting for the total number of genes per quadrant and number of genes in each quadrant on each contig and is therefore particularly well-suited for detecting non-random localised patterns such as secreted gene clustering.

Hypergeometric tests (HgT) were used to evaluate enrichment for functionally characterised genes amongst quadrants defined by flanking intergenic regions. To compare *p*-values across real genes, a test was performed in which the number of genes encoding secretion signals was calculated for each of 4,774 eukaryotic nuclear genomes, and that number of genes were randomly selected. Those random genes (the same number as secreted proteins) then had their intergenic distances compared with the remaining genes using HgT. The process was repeated 1000 times per genome (i.e., 1,000 bootstraps) and visualised in **Fig. 2D**.

**Repeat enrichment flanking genes encoding secreted signals**

To assess whether certain repeat types were significantly enriched or depleted flanking genes with or without predicted secretion signals, repeat counts were compared using *t*-tests and Wilcoxon rank-sum tests in R. For each repeat type (e.g. DNA_Fungi, LINE_Metazoa, etc.), we compared distributions of per-assembly repeat counts between categories. To control for multiple testing, we applied Benjamini-Hochberg correction to both *p*-value sets and considered adjusted *p*-values (FDR) < 0.01 to be statistically significant. We further assessed directionality of differences by comparing group means.

**Supplementary Results**

Phylogenetic trees were constructed from core eukaryotic genes from all genome assemblies passing CEG analysis, and a minimum gene count of 182 illustrates broad taxonomic representation across fungi, metazoa, plants, and other eukaryotes (**Figs. 1A; S1-S8**). Across all eukaryotic genome assemblies analysed, Durum wheat (*Triticum turgidum* subsp. *durum*) has the greatest number of genes with secreted signals (*n* = 18,752/190,470; 9.8%) followed by bread wheat **(***Triticum aestivum;* 17,686/135,934; 13%). The species with the largest percent of genes with secretion signals is the amphibian pathogen *Batrachochytrium salamandrivorans* (*n* = 2,353/11,454; 20.54%) which has a large expanded secretome including several protease families thought to break down host skin and extracellular matrix ^13^. The species with the 2^nd^ greatest percent of genes with secretion signals (3,887/19,128; 20.32%) is the sleeping chironomid (*Polypedilum vanderplanki*), which is a non-biting midge that exhibits extreme tolerance to complete desiccation and secretes an abundance of trehalose from their fat body into the hemolymph during desiccation ^14^.

While the percent of genes encoding secreted proteins are not normally distributed (Shapiro-Wilk test, using normal distribution; right-tailed), as visualised in the histogram in **Fig. 1B**, the observed effect size is very small (KS - D = 0.03896), and therefore comparisons using *t*-tests are appropriate. Two-tailed *t*-tests (assuming unequal variance) suggests differences between the percent of genes with secretion signals compared with genes without secretion signals across each kingdom (for animal, fungi, plant, other*, p* = 0.01, 3.04E^-5^, 7.32E^-18^ and 1.27E^-5^, respectively). Assuming equal variance for the *t-tests* slightly decreases *p*-values (i.e., increases significance). Among the top 100 species with the greatest percent of genes that encode secretion signals, 71 are animal (Hypergeometric test; HgT with upper tail *p*-value = 1.66E^-32^). Conversely, the bottom 100 species comprise 36 animals (also enriched: HgT *p*-value = 6.12E^-6^) and 36 plantae (HgT *p*-value = 7.43E^-18^).

Genes with secretion signals have longer 3’ and 5’ flanking intergenic regions (FIR) than non-secreted proteins among the eukaryotes (**Fig. 2A-B**). Specifically, genes encoding signal peptides had 321 nucleotides (nt) longer 3’ x̃ FIR (1,534 nt > 1,213 nt) and 31 nt longer 5’ x̃ FIR (1,025 nt > 994 nt) than genes without signal peptides based on median values. However, the mean (x̄) FIR for genes encoding secreted proteins (3’ x̄ = 9,729 nt, 3’ standard deviation/σ = 53 Kb, 5’ x̄ = 7,660 nt, σ = 45 Kb) were slightly shorter than non-secreted proteins (3’ x̄ = 9,914 nt, 3’ σ = 50 Kb, 5’ x̄ = 8,314 nt, 5’ σ = 46 Kb), possibly owing to large outliers (as suggested by the σ). Gene outliers included the human encoded *VAMP7* gene that resides within the pseudo-autosomal 2 region of the Y-chromosome and is flanked by *PRYP3* that is 38.6 Mb downstream and the interleukin 9 receptor that is 53.8 Kb upstream (**Fig. S7**). The UCSC Genome Browser revealed several closer features that were absent from the downloaded annotation file, including the closest *CCNQP2* pseudogene 2 that is 35.7 Mb downstream, which is still very distant, and more so than the 2^nd^ longest detected at 28.2 Mb. Therefore, the FIR length between *VAMP7* and *CCNQP2* is the greatest detected among the eukaryotes and contributes to the very long mean of certain genes (in this case without a secretion signal). Using median values (that are not as impacted by such outliers as the mean is) demonstrates longer FIR for genes encoding secretion signals.

Comparing the FIR for all eukaryotic genes with secretion signals (*n* = 5,213,813) against all other genes (*n* = 68,128,352) revealed a highly significant enrichment for genes with secretion signals with > log_10_ median 3’ FIR and > log_10_ median 5’ FIR (visualised as Quadrant Upper-Right; Q_UR_; Hypergeometric test; HgT *p*-value < 2.22e^-308^, which is the smallest non-zero normalized floating-point number reported by R), as well as those with < log_10_ median 5’ FIR (Quadrant Lower-Right; Q_UL_; HgT *p*-value < 2.22e^-308^) (**Figs. 2C; S10**). Indeed, few genes encoding secretion signals were identified with < log_10_ median 3’ FIR and < log_10_ median 5’ FIR (Quadrant Lower-Left; Q_LL_; HgT *p*-value = 1) or > log_10_ median 5’ FIR (Quadrant Upper-Left; Q_UL_; HgT *p*-value = 1). Most (58.4%) individual genome assemblies have a greater number of genes with secretion signals in Q_UR_ than Q_LL_. Computing the significance of genes with secretion signals with long FIR for individual species demonstrated that the majority of species (*n* = 2,483; 53%) had a significant abundance of genes encoding secretion signals in Q_UR_ (HgT α = 0.01, ∴ Bonferroni correction where *n* = 4,694 = *p*-value < 2.13E^-6^) and 1,384 (29%) of species in Q_LR_ (**Fig. 2C, Dataset S1**). Conversely, only 3 species (< 0.1%) and 31 species (0.7%) had significant HgT for genes with secretion signals in Q_UL_ and Q_LL_, respectively. No correlation was identified between the significance of enrichment for genes encoding secreted proteins in each quadrant and the number or percent of genes encoding secreted proteins (**Fig. S11**), suggesting their association with longer FIRs are not a direct result of their abundance or copy number in any given genome.

To test the accuracy of the enrichment protocol, the process was repeated by counting the number of genes with secretion signals in each genome assembly, then randomly selecting that number of genes irrespective of the protein they encode, followed by performing the same enrichment tests. This process was performed on every Eukaryote genome assembly 1000 times (i.e., bootstrap analysis) (**Fig. 2D**). The results from this random allocation bootstrap experiment revealed that no quadrant achieved significant enrichment < *p*-value of 1E^-7^, contrasting the enrichment identified from genes encoding secretion signals that reached *p* ≤ 2.22e^-308^ (only *p*-values accurately reported by R shown in **Fig. 2C**).

To determine how often consecutive/flanking genes are in the same quadrants across each chromosome or contig, a time-discreet custom Markov model was used to identify consecutive genes in a quadrant compared with random chance alone, considering the total number in each quadrant within each contig. This approach identified 615 regions (284 Q_UR_ and 331 Q_LL_) containing between 14 and 217 consecutive genes (based on the Bonferroni correction adjusted *p*-value using the number of assemblies: < 1.97E^-10^; **Dataset S2**). Chromosome 3 of *Solanum tuberosum* (potato) has the most significant (*p* = 2.66E^-96^) number of consecutive genes (*n* = 213/3,815) in any given quadrant (in this case, Q_LL_). The proceeding 13 most significant regions were also from Q_LL_, including *Olea europaea subsp. europaea* (common olive; *p* = 4.29E^-76^), *Trichonephila clavipes* (golden silk spider; *p* = 2.23E^-55^), and the herb *Senna tora* (sickle senna; *p* = 1.35E^-53^).

To assess the relationship between repetitive elements and genes encoding secretion signals, repeats were predicted in a subset of 393/4,694 genome assemblies (8.4%), selected alphanumerically by GCA accession (**Datasets S3-S4**). This subset included 227 fungal genomes (7.2%), 55 metazoan (6.4%), 29 plant (9.6%), and 82 from other eukaryotic groups (20.4%). Comparing the 8.4% of assemblies to the full set yielded t-tests of *p* > 0.01 for gene count, gene count with secretion signals, the percent of genes with secretion signals and the N50, while p = 0.0026 for genome length. When performing the tests of each of the kingdoms, we found the subset of assemblies were different for fungi (p < 0.01), suggesting that the repeat-calling results may not be as generalisable to the full fungal kingdom as the others (**Tables S1**).

Across the 8.4% of genome assemblies, repeat content flanking genes with and without secretion signals was quantified and normalised per gene to account for variation in gene counts. Consistently across kingdoms, repeat elements were significantly more numerous flanking genes with secretion signals, with the strongest enrichments observed in fungi (**Fig. 3**). *t*-tests revealed that simple repeats, low-complexity elements, and repeats of unknown classification were particularly enriched near genes with secretion signals in fungal genomes (adjusted *p* < 0.001) (**Tables S2-S3**).

Although genes encoding secretion signals are flanked by a greater number of repeat elements, the individual repeats surrounding genes lacking secretion signals are consistently longer (**Figs S13-S17; Tables S4-S6**). This pattern holds across most repeat families and kingdoms, regardless of whether repeat lengths are normalised by the number of flanking genes. Statistical comparisons using *t*-tests confirmed significantly longer repeats around genes lacking secretion signals across nearly all repeat categories.

**Supplementary Figures**

**Figure S1.** The percentage of core eukaryotic gene (CEG) coverage across 800 eukaryotic genome assemblies, based on alignments to a curated set of 225 core eukaryotic genes. Coverage represents the proportion of annotated protein-coding genes in each assembly that have ≥ 70% alignment to a CEG. Assemblies shown in red were excluded from downstream analyses due to having < 50% of protein-coding genes meeting this threshold.

**Figure S2.** Continuation of CEG coverage barplots for a second batch of 800 high-quality eukaryotic genome assemblies.

**Figure S3.** Continuation of CEG coverage barplots for a third batch of 800 high-quality eukaryotic genome assemblies.

**Figure S4.** Continuation of CEG coverage barplots for a fourth batch of 800 high-quality eukaryotic genome assemblies.

**Figure S5.** Continuation of CEG coverage barplots for a fifth batch of 800 high-quality genome assemblies.

**Figure S6.** Continuation of CEG coverage barplots for a final sixth batch of 773 high-quality genome assemblies.

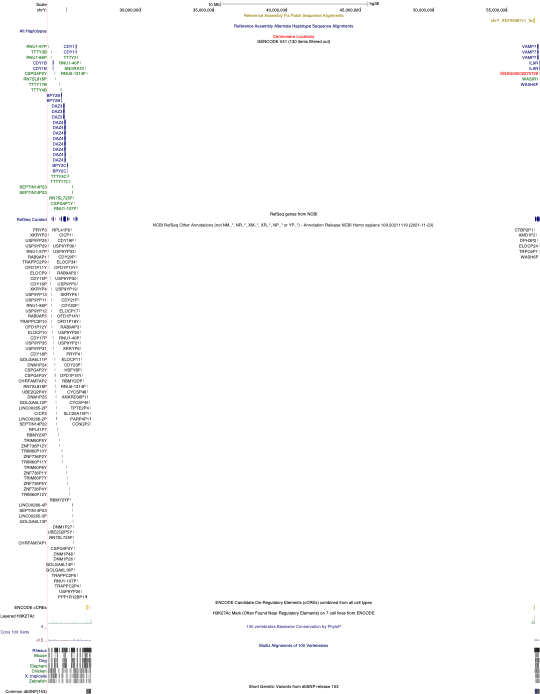

**Figure S7.** Screenshot of the UCSC Genome Browser on Human (GRCh38/hg38) of chrY:20,984,916-57,227,415, showing the longest flanking intergenic region detected in the eukaryote annotation (VAMP7 and PRYP3), which is 38.6 Mb apart. The browser shows several additional features that were not described in the Genbank GFF which are closer albeit still very distant, such as the CCNQP2 pseudogene 2 at positions 26626323-26627368, and therefore making the intergenic region 35.7 Mb from VAMP7.

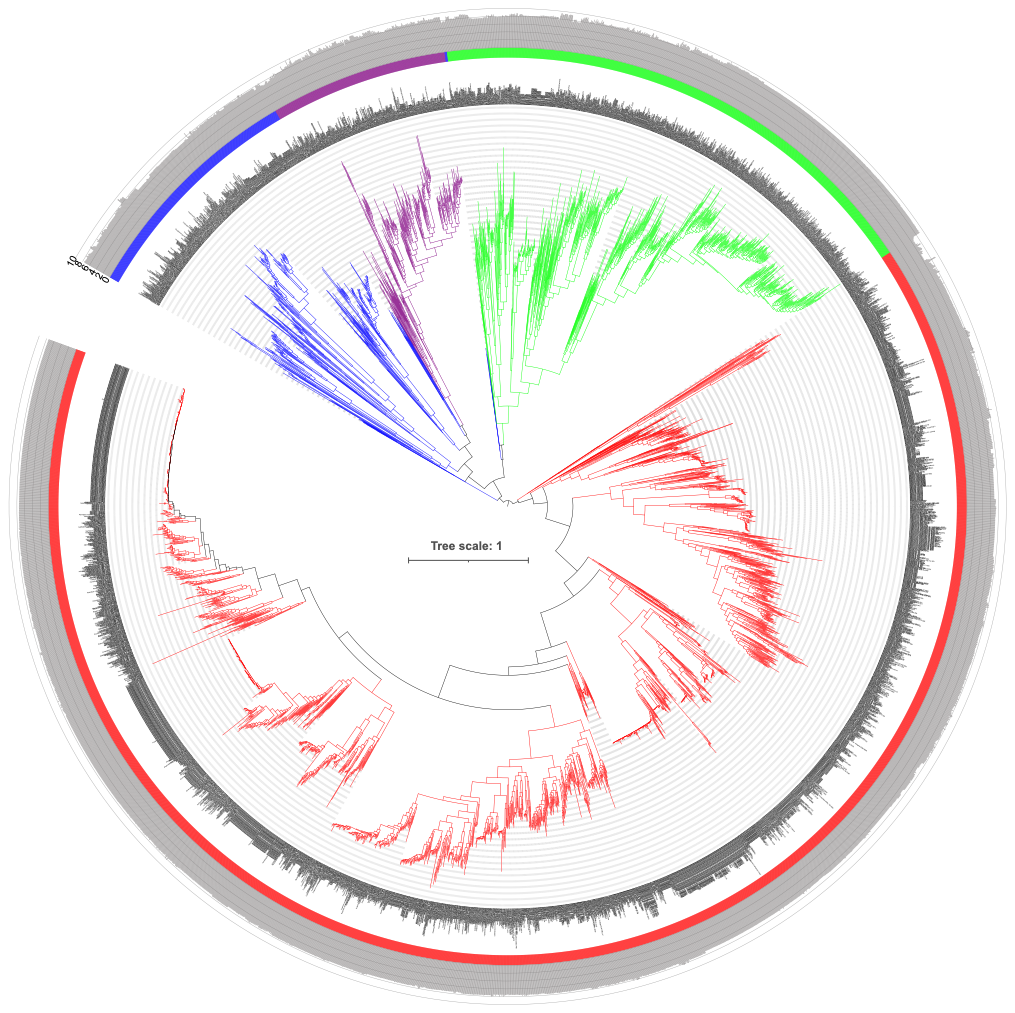

**Figure S8.** Phylogenetic tree of 4,694 eukaryotic species, constructed from multiple sequence alignments of 125 core eukaryotic genes (alignment length (amino acids and gaps) = 714,761 per taxa) and inferred using a midpoint-rooted FastTree. Surrounding the tree are the GCA genome assembly labels, and the genome lengths in grey, shown as a log_10_ scale. Green = animal, red = fungi, purple = plant, blue = other.

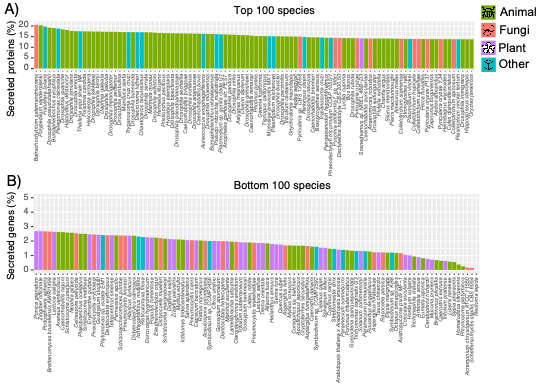

**Figure S9.** Percentage of genes encoding secretion signals across 4,774 eukaryotic genome assemblies, ordered from highest to lowest. (**A**) the kingdom composition for the top 100 and (**B**) bottom 100 genomes based on secreted protein percentages. Green = animal, red = fungi, purple = plant, blue = other.

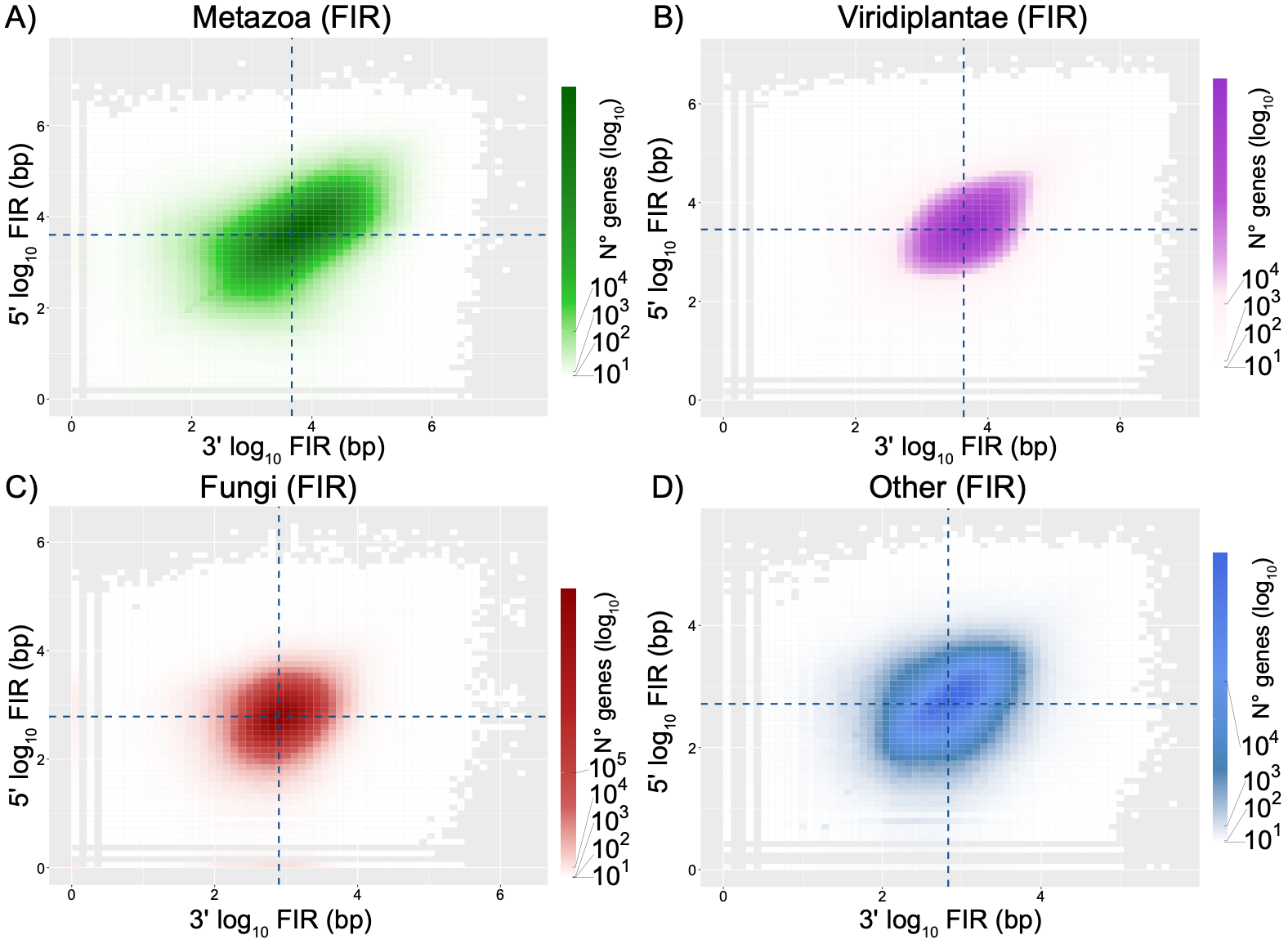

**Figure S10.** The flanking intergenic region (FIR) lengths for each eukaryotic kingdom. Dotted lines indicate the median FIR values, which define four quadrants: upper left (UL), upper right (UR), lower left (LL) and lower right (LR).

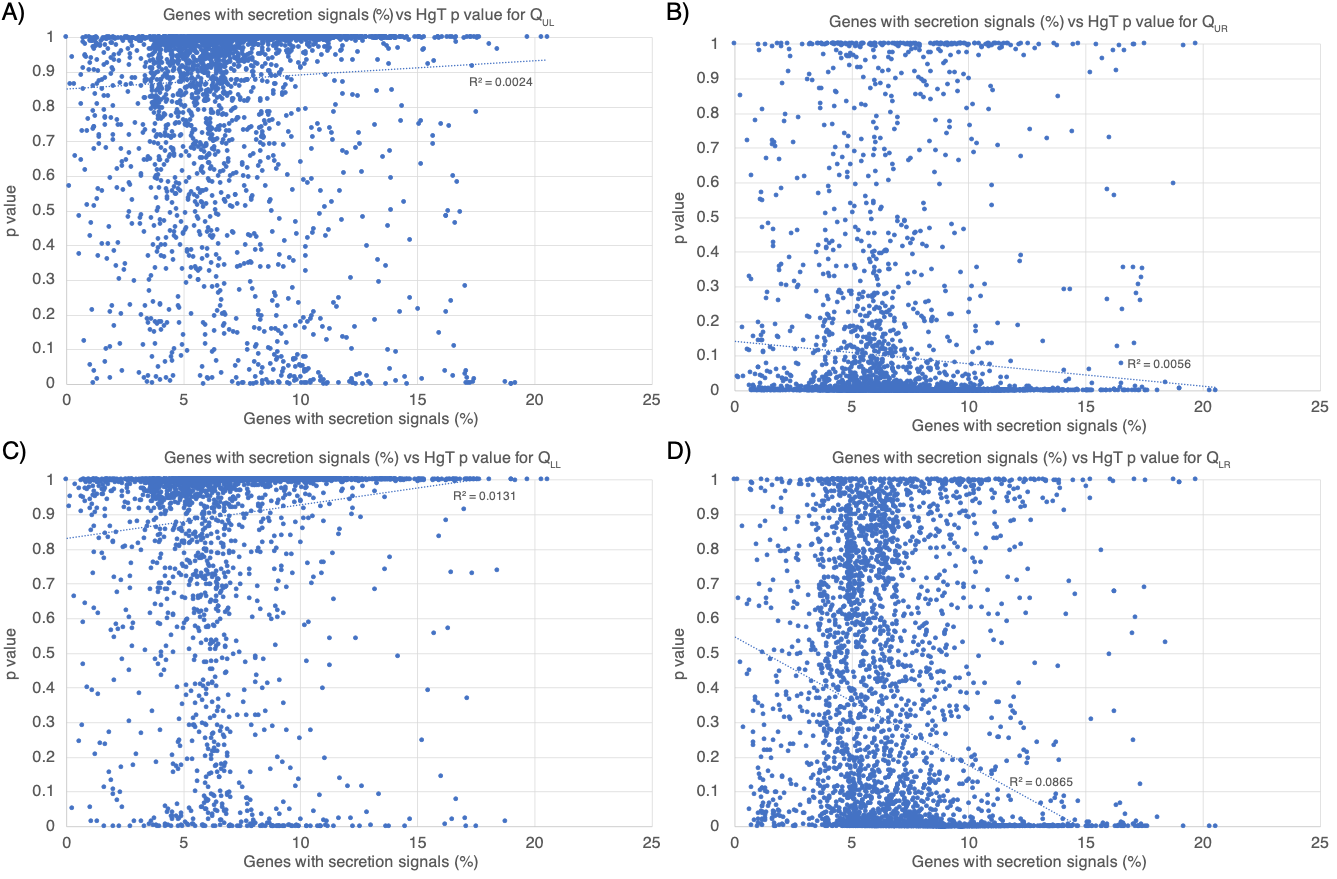

**Figure S11.** Percentage of genes with secretion signals were analysed for enrichment in each of the 4 quadrants across all 4,694 eukaryotic species. Quadrants were defined by the median flanking intergenic region (FIR) lengths: upper left (UL), upper right (UR), lower left (LL) and lower right (LR). Enrichment was testing using hypergeometric tests (HgT), with *p*-values shown as the y-axis. No clear correlation was identified between the significance of enrichment for genes encoding secreted proteins in each quadrant. R^2^ value and line of best fit is shown for each quadrant.

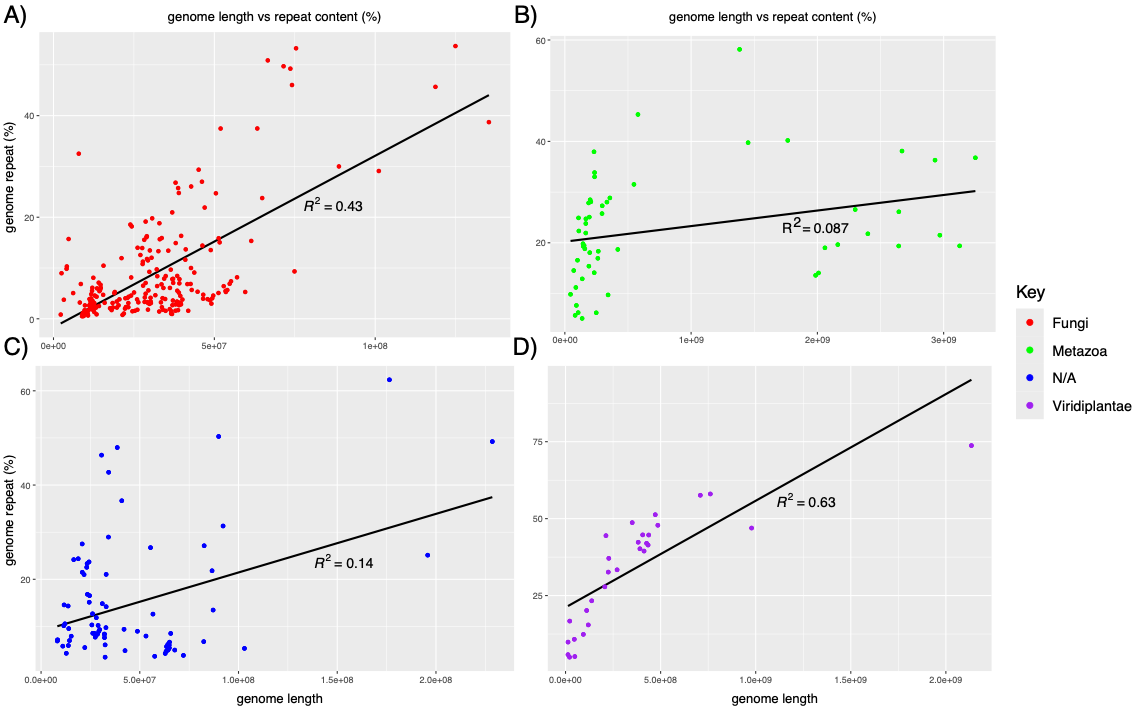

**Figure S12.** Genome length vs repeat content (as a percent of genome length) for (**A**) Fungi, (**B**) Metazoa, (**C**) Other and (**D**) Plants. Repeats were calculated for only the first 393/4,694 assemblies (ordered alphanumerically by GCA number), representing

8.4% of all available eukaryotic genome assemblies including 227 Fungi (7.2%), 55 Metazoa (6.4%), 29 Plants (9.6%), and 82 Other (20.4%) R^2^ value and line of best fit is shown for each.

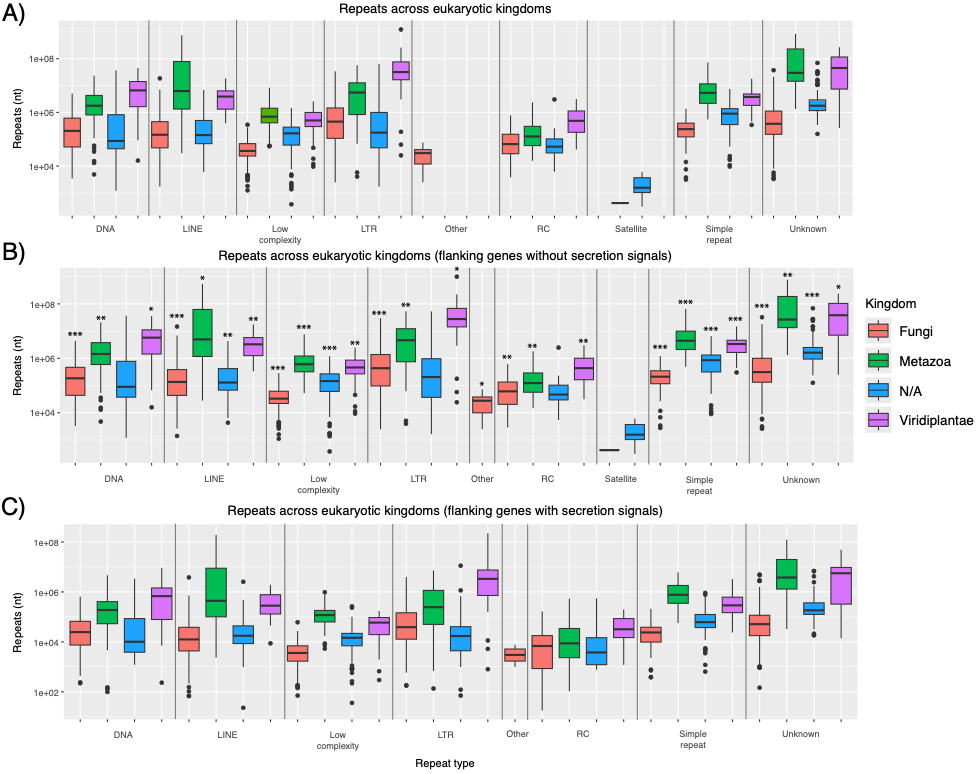

**Figure S13.** The distribution of repeat lengths (in nucleotides) across the eukaryotic kingdoms for each repeat superfamily. Colours represent taxonomic groups: red = fungi, green = animals, purple = plants, blue = other. (**A**) Total repeat lengths for all repeats, regardless of gene context, across eukaryotic kingdoms. (**B**) Repeat lengths for elements flanking non-genes lacking secretion signals. (**C**) Repeat lengths for elements flanking genes encoding secretion signals. Significance testing between Panels B and C was performed using unpaired two-sided t-tests, with significance levels indicated by asterisks above Panel B *** = *p* < 0.001, ** = *p* < 0.01, * = *p* < 0.05.

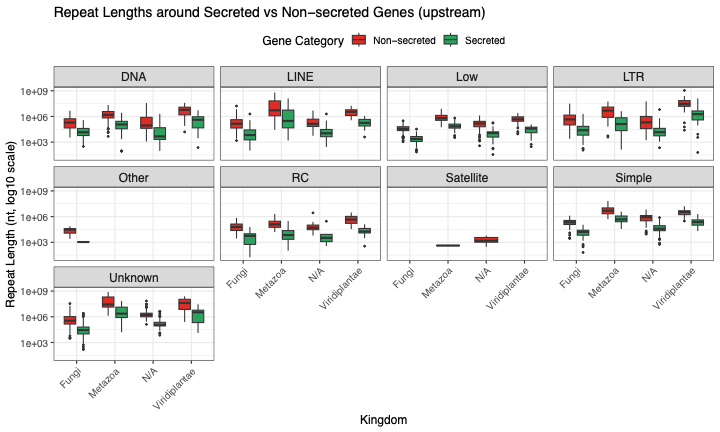

**Figure S14.** The distribution of repeat lengths (in nucleotides) upstream of genes with or without secretion signals, across the eukaryotic kingdoms for each repeat superfamily.

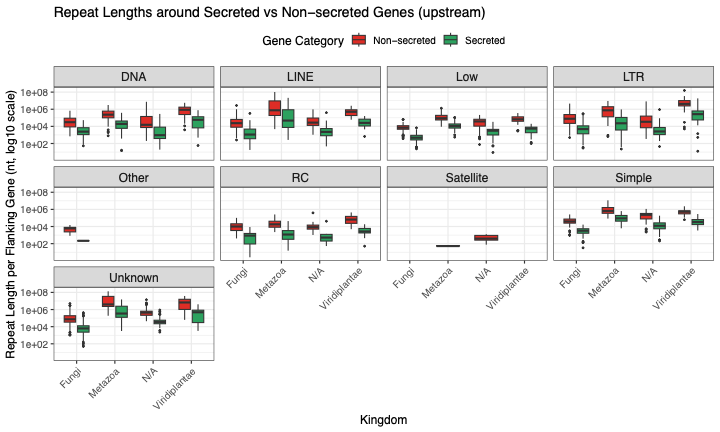

**Figure S15.** The distribution of repeat lengths (in nucleotides) upstream of genes with or without secretion signals, normalised by the number of genes that with or without secretion signals, across the eukaryotic kingdoms for each repeat superfamily.

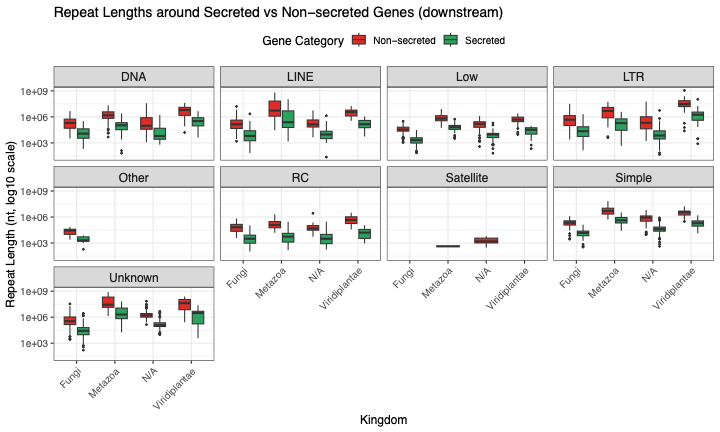

**Figure S16.** The distribution of repeat lengths (in nucleotides) downstream of genes with or without secretion signals, across the eukaryotic kingdoms for each repeat superfamily.

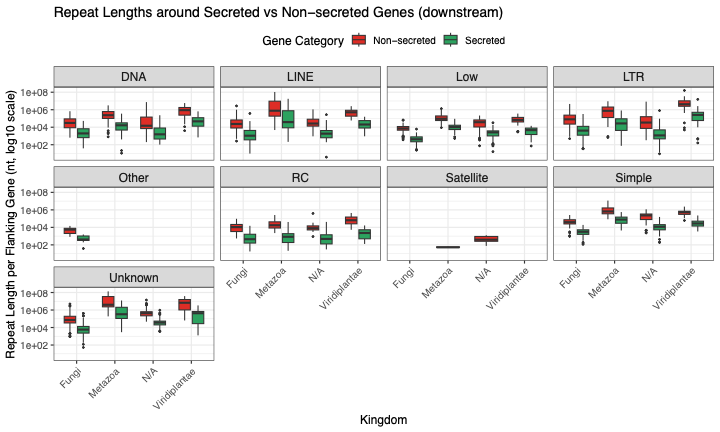

**Figure S17.** The distribution of repeat lengths (in nucleotides) downstream of genes with or without secretion signals, normalised by the number of genes that with or without secretion signals, across the eukaryotic kingdoms for each repeat superfamily.

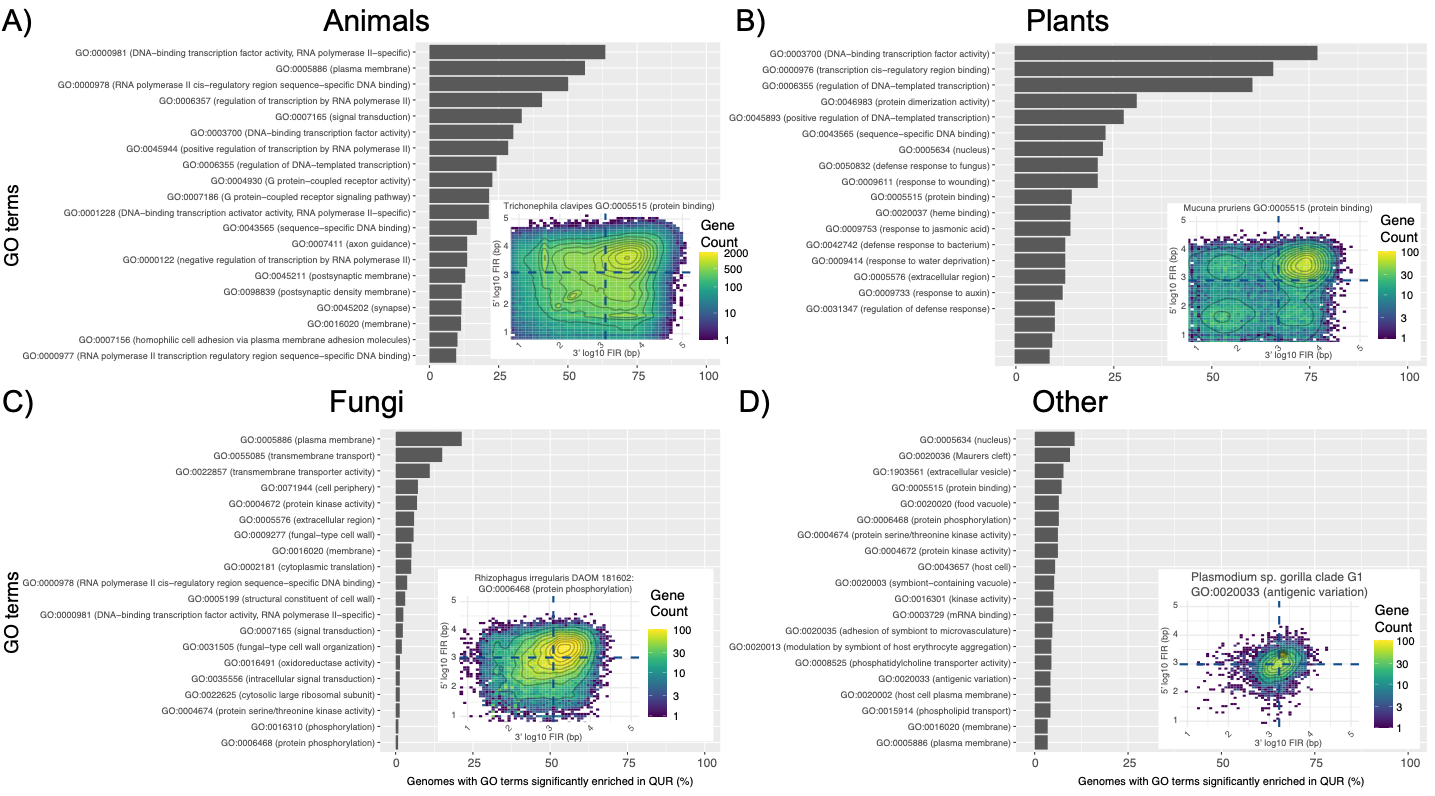

**Figure S18.** The most frequently enriched GO-terms in the Q_UR_ for each of the four kingdoms (A = Animals, B = Plants, C = Fungi, D = Other). The most significantly enriched terms had their flanking intergenic region (FIR) plotted for illustrative purposes, as shown in the bottom right of each barchart.

**Supplementary Tables**

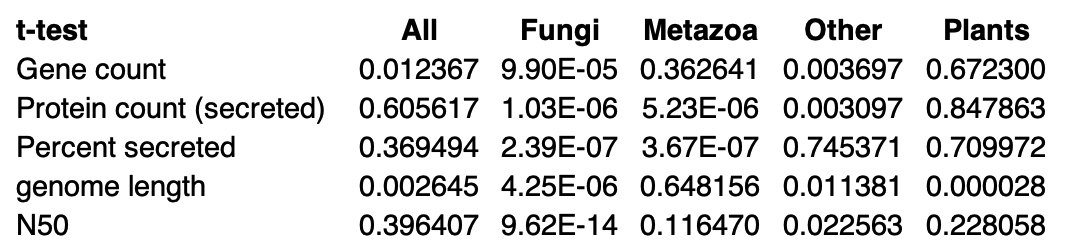

**Table S1.** Welch unequal-variance *t*-tests were used to compare various assembly metrics of the 8.4% of genome assemblies used for repeat calling, compared with the values in the full dataset. p-values are shown for comparisons across all assemblies and limiting the comparisons to individual kingdoms. While the 8.4% of ‘All’, Other and Plants appeared to be a good representative of the full set based on these tests, fungi and to a lesser extent metazoa had significant differences.

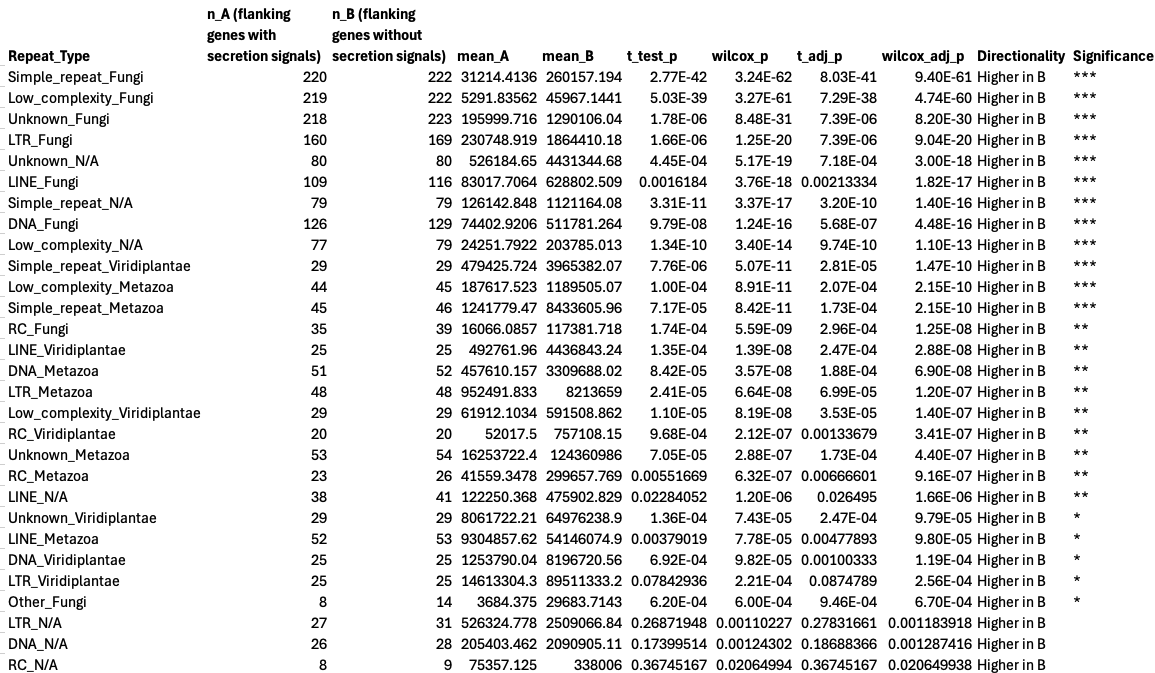

**Table S2.** *t*-tests were used to compare the number of repeats that were flanking genes with secretion signals, compared to genes without secretion signals. Significance levels indicated by asterisks where *** = *p* < 0.001, ** = *p* < 0.01, * = *p* < 0.05.

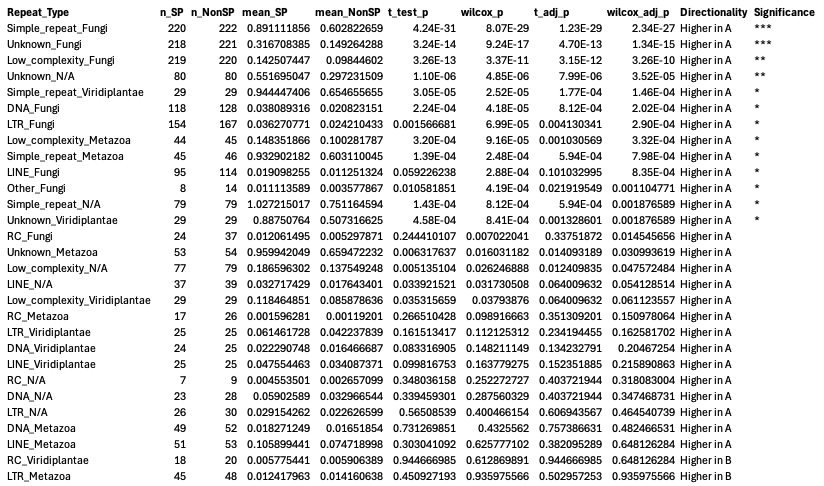

**Table S2.** Due to different numbers of genes with or without secretion signals, the t-tests in Table S1 were re-calculated, where the numbers of genes flanked by a repeat were divided (normalised) by the total numbers of those genes (i.e., with or without a secretion signal). Significance levels indicated by asterisks where *** = *p* < 0.001, ** = *p* < 0.01, * = *p* < 0.05.

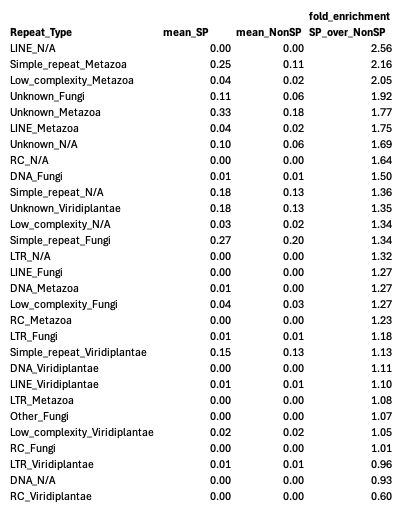

**Table S3.** The fold enrichment of genes with secretion signals flanking repeats (normalised by number of genes with secretion signals) compared to the numbers of genes without secretion signals (normalised by number of genes without secretion signals).

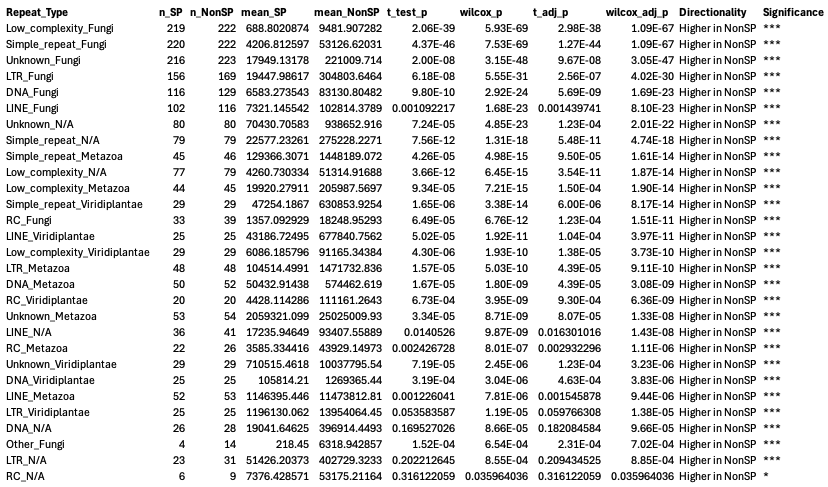

**Table S4.** *t*-tests were used to compare the number of repeats (nucleotides) upstream of genes with secretion signals, compared to genes without secretion signals. Significance levels indicated by asterisks where *** = *p* < 0.001, ** = *p* < 0.01, * = *p* < 0.05.

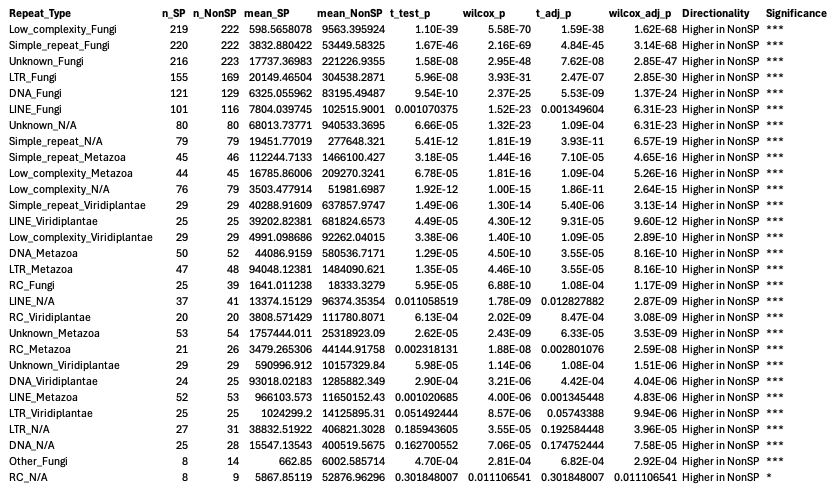

**Table S5.** *t*-tests were used to compare the number of repeats (nucleotides) downstream of genes with secretion signals, compared to genes without secretion signals. Significance levels indicated by asterisks where *** = *p* < 0.001, ** = *p* < 0.01, * = *p* < 0.05.

**Supplementary Dataset Legends**

**Dataset S1.** The nexus file generated from 125 core eukaryotic genes (CEGs) identified from 4,694 genome assemblies with blast v2.2.24 ^8^. Multiple sequence alignments were made using MUSCLE v3.8.31 ^9^, refined with trimAl v1.4.1 ^10^ and a phylogenetic tree was constructed with FastTree v2.1.11 ^11^.

**Dataset S2.** The percent of genes with secretion signals in each of the 4 quadrants for all genome assemblies (a separate tab for each quadrant). The dataset includes the assembly, assembly name, the kingdom, phylum, species, percentage of genes with secretion signals (secreted %), and the hypergeometric test *p*-value for having that number or a greater number (upper tail) of genes with secretion signals in that quadrant. The number of genes and genes encoding secretion signals are given in a fifth tab.

**Dataset S3.** The results from the custom time-discrete Markov model, accounting for the number of genes in each quadrant on each contig and using a Bonferroni-adjusted significance threshold (*p* < 1.97 x 10^-10^). The dataset includes the assembly, the contig, the region saved (genomic coordinates), which quadrant (1 = Q_UL_, 2 = Q_UR_, 3 = Q_LR_, and 4 = Q_LL_), the number of consecutive genes in that region (Consec_count), the total count of genes on that contig (Total_count), the probability of having a gene in that quadrant (Prob_quadrant), and the cumulative probability of having that number of consecutive genes in that quadrant (Cumulative_probability_of_consecutive_genes).

**Dataset S4.** All genome assemblies that were repeat called. Those highlighted in red had no repeats identified. The dataset includes the assembly, assembly name, Taxonomy ID (TaxID), kingdom and species name.

**Dataset S5.** Information about the repeats called, including genome information (summary 1/tab 1), repeat info for subfamilies (summary 2/tab 2), repeat info for repeat families (summary 3/tab 3), and a summary of assemblies (completed/tab 4).

**Dataset S6.** Go-term enrichment in genes encoding secretion signals compared with genes not encoding secretion signals. Enrichments were made separately in each genome assembly, and then in this dataset, the tally of genomes those terms were enriched in were recorded, where Tab 1 = All kingdoms/genome assemblies, Tab 2 = Fungi only, Tab 3 = Animals only, Tab 4 = Plants only and Tab 5 = ‘Other’ kingdom only.

**Dataset S7.** The individual genome-based data that is summarised in **Dataset S6**, showing the genome assembly accession and q-value for each enriched GO-term among genes encoding secretion signals compared with genes not encoding secretion signals.

**Dataset S8.** All GO-terms in each genome assembly were analysed for enrichment in each of the four quadrants. The results are separated into tabs reflecting the quadrant analysed and the GO-terms identified. The genome from which the enrichment was identified is also recorded. Enrichment is considered where q-values < 0.00001. The generic GO:0003674 (molecular_function) was excluded from the results.

**Dataset S9.** A summary of **Dataset S8**, where the number of genome assemblies that are enriched for GO-terms in each quadrant are recorded. Enrichment is considered where q-values < 0.00001. The generic GO:0003674 (molecular_function) was excluded from the results.
